# Ancestral intronic splicing regulatory elements in the SCNα gene family

**DOI:** 10.1101/2025.08.16.670673

**Authors:** Ekaterina Chernyavskaya, Margarita Vorobeva, Sergei A. Spirin, Dmitry A. Skvortsov, Dmitri D. Pervouchine

## Abstract

SCNα genes encode components of voltage-gated sodium channels that are crucial for generating electrical signals. Humans have ten paralogous SCNα genes, some of which contain duplicated mutually exclusive exons 5a and 5b. In reconstructing their evolutionary history, we found multiple unannotated copies of exon 5 in distant species and showed that exon 5 duplication goes back to a common ancestor of the SCNα gene family. We characterized splicing patterns of exons 5a and 5b across tissues, tumors, and developmental stages, and demonstrated that the nonsense mediated decay (NMD) system is not the major factor contributing to their mutually exclusive choice. Comparison of *SCN2A*, *SCN3A*, *SCN5A*, and *SCN9A* intronic nucleotide sequences revealed multiple Rbfox2 binding sites and two highly conserved intronic splicing regulatory elements (ISRE) that are shared between paralogs. Minigene mutagenesis and blockage by antisense oligonucleotides showed that the formation of RNA structure between ISRE promotes exon 5b skipping in *SCN9A*. The inclusion of exon 5b is also suppressed in siRNA-mediated knockdown of Rbfox2, which makes the collective action of RNA structure and Rbfox2 compatible with the model of a structural RNA bridge. ISRE sequences are conserved from human to elephant shark and may represent an ancient, evolutionarily conserved regulatory mechanism. Our results demonstrate the power of comparative sequences analysis in application to paralogs for elucidating splicing regulatory programs.

## Introduction

Voltage-gated sodium channels are crucial for generation and propagation of action potentials in excitable cells including neurons, muscle cells, and some glial cells. The α-subunits of these channels, encoded by the SCNα gene family, form the principal ion-conducting pore [1]. Each α-subunit consists of four homologous domains, with each domain containing six transmembrane segments (S1–S6). The voltage-sensing module includes segments S1–S4 and undergoes conformational changes in response to membrane potential alterations, particularly through the movement of charged residues in the S4 segment, while segments S5–S6 form the sodium-selective pore [2, 3, 4].

The human SCNα family comprises ten paralogs (*SCN1A–SCN11A*, where *SCN6A* and *SCN7A* is the same gene) which exhibit distinct tissue-specific expression patterns [1]. For instance, *SCN1A–SCN3A* and *SCN9A* are predominantly expressed in the nervous system, whereas *SCN4A* is found to be activated in skeletal muscle [5, 6]. A notable feature of *SCN1–3A*, *SCN5A*, *SCN8A* and *SCN9A* is the presence of splice isoforms with mutually exclusive exons 5a and 5b (MXE), the expression of which is tightly regulated during embryonic development [7, 8]. Imbalanced MXE isoform composition causes altered sodium channel permeability that has been linked to various pathologies, including cardiac-conduction delay, neuropathic pain, and epilepsy [9, 10, 11]. Similar alterations in the relative abundances of splice isoforms of other genes, particularly, voltage-gated calcium channels are involved in a broad group of disorders termed channelopathies [12].

Exons 5a and 5b are both 92-nts-long, hence their coordinated inclusion or skipping would cause a frameshift and generate a transcript isoform with a premature termination codon. Elimination of such transcripts by the nonsense mediated decay (NMD) pathway has been proposed as a plausible mechanism underlying mutually exclusive splicing in clusters with exactly two exons [13]. However, insights from other MXE systems suggest a potential role of RNA structure. A well-characterized example is the *Drosophila Dscam1* gene, in which large clusters of MXEs are controlled by competing RNA structures [14, 15, 16]. It has been proposed that the acquisition of competing RNA structures and MXEs is a natural consequence of tandem genomic duplications which, on the one hand, generate a group of homologous exons and, on the other hand, produce a group of sequences that compete for complementary base pairings with another sequence [17, 18, 19].

Regulatory RNA structures are anecdotally found in paralogues [20]. A notable example is the group of RNA structures in the fibroblast growth factor receptor (FGFR) gene family, which control mutually exclusive splicing of exons IIIb and IIIc, simultaneous inclusion or exclusion of which also can invoke NMD [21]. Complementary base pairings between intronic elements IAS2 and ISAR in the *FGFR2* gene have been conserved from sea urchin to humans [22], while sharing a common sequence with complementary intronic elements ISE-2 and ISAR in its paralog *FGFR1* [23]. In this case, regulatory RNA structures probably had been present in the common ancestor of *FGFR1* and *FGFR2* before gene duplication occurred. However, the example of the BET gene family demonstrates that RNA structures regulating alternative splicing in paralogs can also be acquired independently in the course of convergent evolution [24].

The SCNα genes represent one of the most studied gene families with respect to their function [25, 26], structure [27, 28], evolution [29, 30], role in disease and drug resistance [31, 32, 33]. Their common ancestry and, furthermore, a remarkable similarity of exon 5a and 5b sequences motivated us to examine this family in searching for intronic splicing regulatory elements (ISRE) involved in alternative splicing regulation [34]. In this work, we characterize two ISRE shared by most SCNα genes with exon 5 duplications, which are capable of forming RNA structures that impact splicing. In *SCN9A*, we demonstrate that their complementary base pairing, indeed, controls MXE splicing ratio through an RNA bridge connecting exon 5b with a distal Rbfox2 binding site.

## Results

### Exon 5 duplications in the SCN**α** gene family

The human genome contains 10 paralogous SCNα genes distributed across different chromosomes, with five genes (*SCN1A*, *SCN2A*, *SCN3A*, *SCN7A*, and *SCN9A*) clustered on chromosome 2, three genes (*SCN5A*, *SCN10A*, and *SCN11A*) on chromosome 3, and the remaining two (*SCN8A* and *SCN4A*) on chromosomes 12 and 17, respectively. Genes sharing chromosomal locations arose through tandem duplications following the initial dispersion of ancestral genes to different chromosomes, as evidenced by their tandem genomic arrangement.

All ten human SCNα genes share a common exon structure in their core protein-coding part with minor variations in exon lengths, however some of them contain duplicated exon 5 variants (5a and 5b) showing on average 91% sequence identity (Figure 1A). At that, exons 5a from different genes are more similar to each other than they are to the respective exons 5b from the cognate gene (Figure 1B). This observation indicates that exon 5 duplication must have occurred in the common ancestor prior to gene duplication events that generated the current genomic organization of the SCNα gene family.

**Figure 1:**
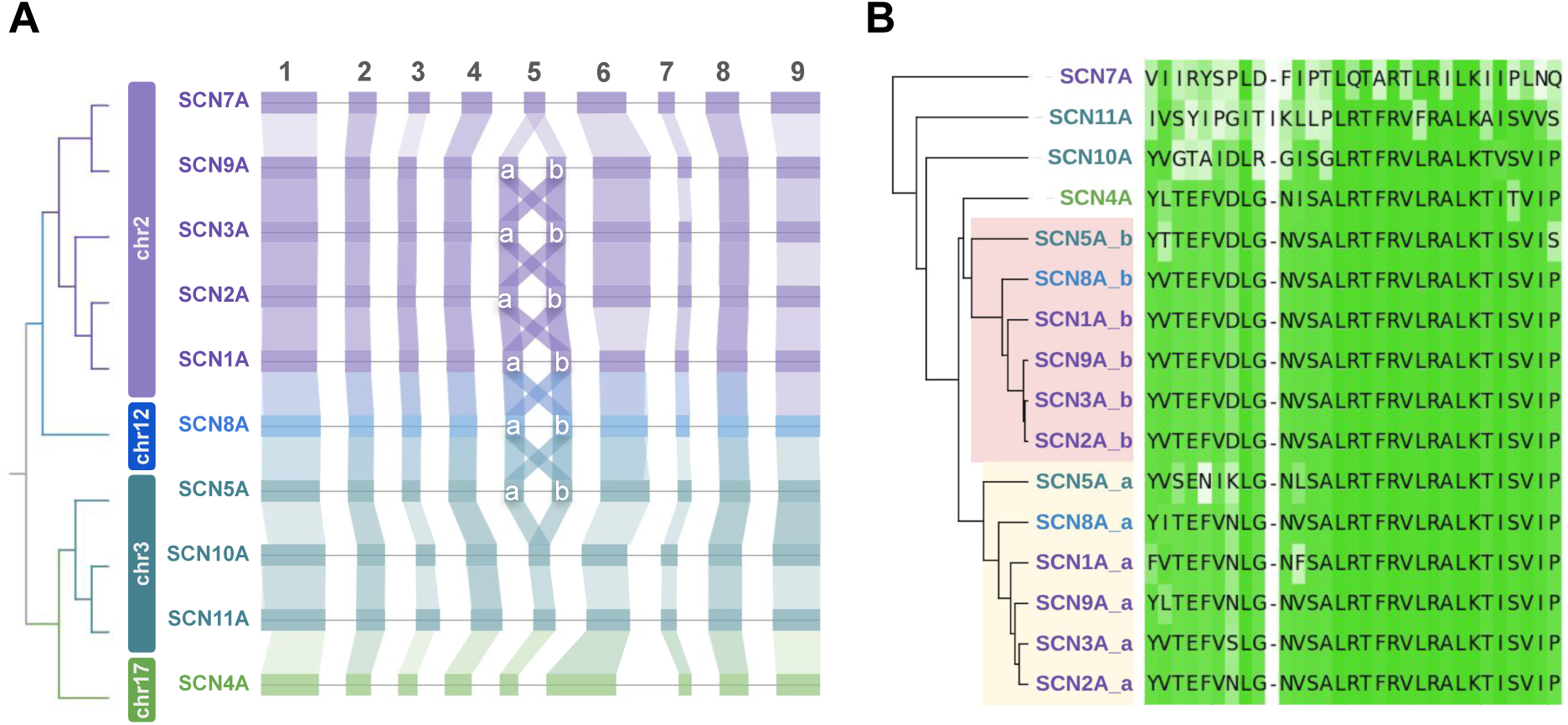
Transcript architecture in the human SCNα gene family **(A)** Exon-exon alignment of the human SCNα genes (first nine exons are shown). Duplications of exon 5 are marked as “a” and “b”. Rectangles on the gene line indicate the annotated exons. The shaded areas connecting exons in different genes illustrate sequence homology of their amino acid sequences, hence the X-like pattern demonstrates that exons 5a and 5b are homologous. **(B)** The alignment of aminoacid sequences of the human SCNα exons 5a and 5b. The intensity of the green background for each amino acid in the alignment reflects its degree of similarity.

To reconstruct the evolutionary history of exon 5 duplications, we searched for orthologs of the ten human SCNα genes (proteins encoded by their major isoforms) in other organisms, with the most distant species being *Aplysia californica* and *Drosophila melanogaster*, and constructed a phylogenetic tree (Figure S1). Additionally, we identified unannotated exon duplications by applying tblastn [35] to the translated amino acid sequences of exon 5 as a query and searching for similarly translating sequences in the adjacent introns (see Methods). We found that exon 5 duplications were widely distributed among jawed vertebrates and included several instances of unannotated duplications (Figure 2A, Table S1). In particular, a previously unknown duplication of exon 5 was observed in *SCN2A* homologs in sheep and tropical clawed frog, as well as in *SCN1A* homologs in zebrafish and elephant shark. As might be expected, exon 5 evolutionary dynamics involved not only acquisitions but also subsequent losses. For instance, exon 5 duplication was absent in the *SCN5A* homolog in the tropical clawed frog. Rodents (mouse, chinese hamster, guinea pig) showed the loss of the duplicated exon in *SCN1A* homolog, while their closest phylogenetic relatives, rabbits and alpine marmots, retained two copies of exon 5.

**Figure 2:**
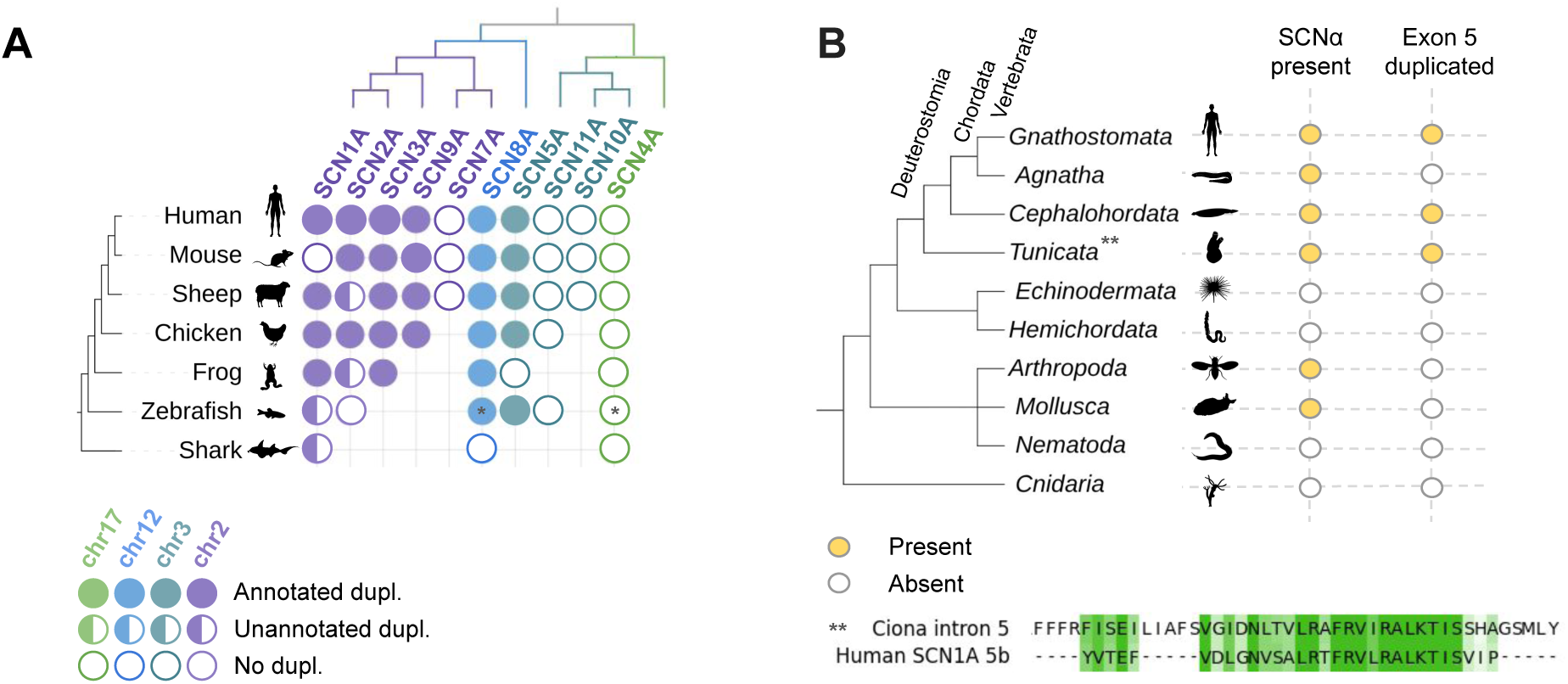
Exon 5 duplications in orthologs of human SCNα genes across species. **(A)** SCNα genes and exon 5 duplications in vertebrates. Asterisk indicates that zebrafish have an additional whole genome duplication, hence *SCN8A* and *SCN4A* orthologs have two duplicated genes, which are not shown here. **(B)** SCNα genes and their exon 5 duplications in animals. Bottom panel: The alignment of translations of the *Cinav1a* gene intron 5 with human *SCN1A* exon 5b, indicating partial exon duplication in *Ciona*.

The phylogenetic trees of SCNα genes in jawless vertebrates and cephalochordata species yielded intriguing results (Figure 2B, the full tree in Figure S1). While no exon 5 duplications were found in jawless vertebrates, potentially due to genome assembly limitations, two out of the five SCNα genes in *B. lanceolatum* contained duplications of the exon homologous to the human exon 5. The gene LOC136428574 (sodium channel protein 1 brain-like) contained an annotated exon 5 duplication, while LOC136422135 (sodium channel protein type 4 subunit alpha B-like) showed an unannotated exon 5 duplication with a remarkable level of sequence identity (Table S1). In tunicates, represented by *Ciona* with three annotated SCNα genes, only a partial duplication of exon 5 in the *Cinav1a* gene was detected, but the level of sequence identity unambiguously supports the occurrence of an ancestral duplication event (Figure 2B, bottom panel). While no duplications of exon 5 homolog were detected in protostomes (arthro-pods and mollusks), duplications of other exons (such as exon 19, which is duplicated in *D. melanogaster*) were still present. Previous studies attributed such duplication patterns to con-vergent evolution in different lineages [36]. Of note, the positions of stop codons that appear as a result of simultaneous skipping or inclusion of the two variants of exons 5 tend to be conserved in most human and related species paralogs (data not shown).

In sum, our findings suggest that exon 5 duplications are characteristic of most human SCNα genes and likely existed in the common ancestor of all chordate species. This duplication may have originated in a common ancestor of chordates, and one of whose genes gave rise to all human SCNα genes. The subsequent evolutionary processes involved multiple independent losses of duplicated exons, particularly in *SCN4A*, *SCN7A*, *SCN10A*, and *SCN11A*.

### Splicing patterns of duplicated exons

In challenging the assumption that exons 5a and 5b are mutually exclusive, we reanalyzed RNA-seq experiments from the GTEx project focusing on six SCNα genes with duplicated exons (*SCN1A*, *SCN2A*, *SCN3A*, *SCN5A*, *SCN8A*, and *SCN9A*). Indeed, these genes exhibit distinct splicing patterns, with *SCN1A*, *SCN2A*, *SCN3A*, and *SCN5A* primarily expressing exon 5b, and *SCN9A* primarily expressing exon 5a (Figure 3A). The 5b/5a transcript ratio shows a moderate degree of variation across tissues, particularly across brain regions, but in *SCN8A* we observed almost a complete reversal of 5b/5a ratio in pituitary and testes, a pattern not observed earlier in any other SCNα gene. Simultaneous inclusion or skipping of both exons were rare, e.g., in *SCN2A* they were both completely skipped in pancreas. However, the expression levels were remarkably low when the two exons were not mutually exclusive, consistent with the degradation of transcripts containing premature stop codons by the NMD pathway. When asking whether exon 5 could be skipped in SCNα genes without duplicated exons, we found no evidence of such skipping in *SCN7A*, *SCN10A*, and *SCN11A*, while in *SCN4A* we observed only a weak skipping of exon 5 in colon and skeletal muscle with the average Ψ < 0.05.

**Figure 3:**
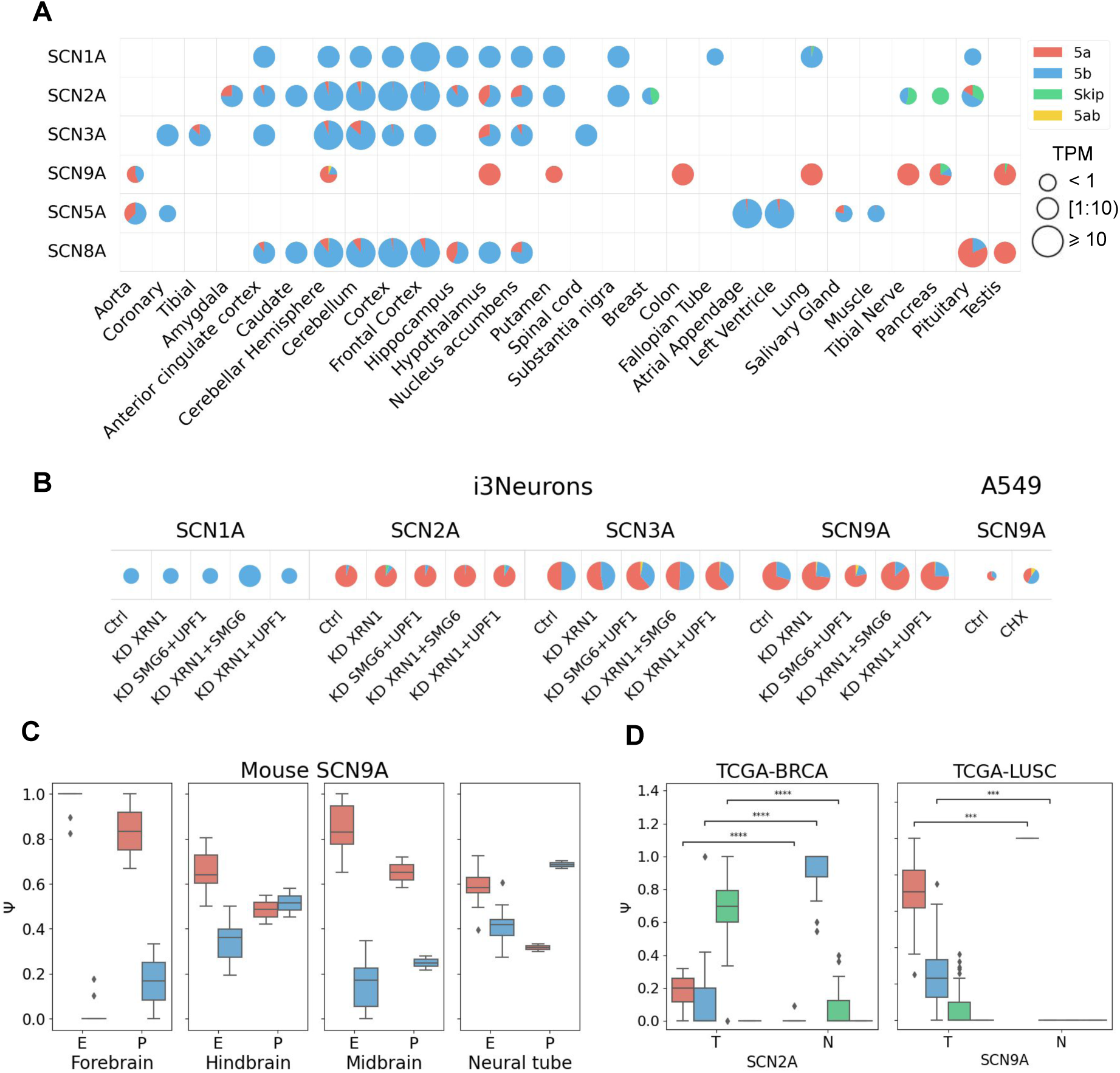
Splicing patterns in SCNα genes. **(A)** Splicing patterns in human tissues. The median Ψ value of each isoform (5a, 5b, 5a5b, and skip) is shown for each tissue. The color legend applies to all panels. The median expression level of a gene is represented by the TPM value (transcripts per million). **(B)** Ψ values of exon 5a and 5b in human cell lines (i3Neurons and A549) in NMD inhibition experiments. KD, Ctrl, and CHX denote the knockdown, the control, and the cycloheximide treatment experiments. **(C)** Developmental dynamics of *SCN9A* exons 5a and 5b in mouse. E — embryonic; P — postnatal. RNA-seq on postnatal stages were made for organs or structures that develop from embryonic ones. The contribution of 5a5b and skip isoforms was zero and is not shown. **(D)** Cancer-specific splicing of exons 5a and 5b in the breast cancer (BRCA) and lung squamous cell carcinoma (LUSC) cohorts from TCGA. Statistically significant differences at the 0.1% significance level are denoted by *** (two-tailed t-test with Bonferroni correction for testing 10 TCGA cohorts).

To examine whether mutually exclusive splicing of exons 5a and 5b is regulated by NMD, we analyzed RNA-seq data from human cell lines, in which the activity of the NMD pathway was suppressed either by the translation inhibitor cycloheximide or by knockdown of key NMD factors. Among SCNα genes with exon 5 duplications, *SCN1A*, *SCN2A*, *SCN3A* and *SCN9A* showed a measurable level of expression in neuronal cells, while in *A549* cells only *SCN9A* was expressed (Figure 3B). We didn’t observe a considerable decline from the mutually exclusive splicing pattern upon NMD inhibition in any of these cases, with the most substantial upregulation of *SCN9A* isoforms containing double exons in *A549* cells (ΔΨ = 0.1). These results indicate that the NMD pathway is not the major factor contributing to mutually exclusive choice of exons 5a and 5b in SCNα genes, thus making RNA structure the next potential suspect.

Previous studies reported significant changes of 5b/5a isoform ratio during development [7, 8]. For example, neonatal *SCN5A* transcripts include exon 5a, whereas adult isoforms predominantly contain exon 5b. To validate this, we reanalyzed RNA-seq experiments from a study of mammalian organ development [37] and mouse embryonic transcriptomes [38]. In all SCNα genes with exon 5 duplications that were expressed at detectable levels, we observed a significant change of the 5b/5a isoform ratio across developmental stages, which was concordant between human and mouse (Figure 3C, Figure S2A). Interestingly, in *SCN9A* we observed a previously undocumented switch in hindbrain and neural tube, where the isoform with exon 5a was dominant in prenatal but not in postnatal samples. On the other hand, the 5b/5a iso-form ratio in postnatal neural tube resembled that of *SCN2A* and *SCN3A* (Figure S2B). No such postnatal switching occurred in the forebrain or midbrain.

Growing evidence indicates that altered SCNα isoform expression is implicated in disease [39, 40, 41]. To assess splicing patterns of SCNα genes in tumors, we reanalyzed RNA-seq experiments from TCGA and found that tumors exhibited a broad spectrum of 5b/5a splicing aberrations: some cases showed complete exon switching, others co-occurred with dual exon skipping, and a subset displayed the loss of tissue specificity (Figure S3). Striking examples in-clude reduced exon 5b inclusion in *SCN2A* (ESCA — esophageal carcinoma; BRCA — breast cancer) and reduced exon 5a inclusion in *SCN9A* (LUAD — lung adenocarcinoma; LUSC — lung squamous cell carcinoma). In BRCA, an elevated fraction of *SCN2A* transcripts lacking both exons was observed suggesting production of nonfunctional mRNA. A similar upregula-tion was also found for *SCN9A* in LUSC (Figure 3D). While these findings demonstrate tumor-associated disruption of mutually exclusive splicing in SCNα genes, they provide no evidence of direct implication of these genes in cancer pathogenesis, being merely a manifestation of global alterations of splicing regulatory programs and NMD pathway efficiency in tumors.

### Common intronic splicing regulatory elements in human SCN**α** genes

The mutually exclusive choice of exons 5a and 5b in SCNα genes hints at the existence of a common regulatory mechanism. The NMD pathway is not the major factor contributing to this mechanism (Figure 3B), and the examples of *FGFR1* and *FGFR2* genes indicate the involve-ment of RNA structure-based splicing regulation. Since *FGFR1* and *FGFR2* share complemen-tary motifs inherited from the common ancestor [23], we examined the introns of SCNα genes for presence of similar common regulatory sequences.

To do this, we constructed a multiple sequence alignment of intronic nucleotide sequences flanking exons 5a and 5b in the six human SCNα genes with exon 5 duplications (*SCN1A*, *SCN2A*, *SCN3A*, *SCN5A*, *SCN8A*, and *SCN9A*). The introns of *SCN1A* and *SCN8A* were too divergent, hence they were analyzed separately. The introns upstream of exon 5a in the re-maining four genes showed almost no similarity besides the expected homology at splice site sequences and polypyrimidine tracts; the intervening introns between exons 5a and 5b were remarkably similar; while the introns downstream of exon 5b contained a pair of ISRE, termed ISRE1 and ISRE2, which shared nearly the same sequence and were complementary to each other (Figure 4, see also Figures S4 and S5 for detailed alignments). According to phastCons score distributions (Figure 5), both regions were evolutionarily conserved across 100 verte-brates, with ISRE1 being located 10–15 nts downstream of exon 5b, and ISRE2 positioned 89–533 nts downstream.

**Figure 4:**
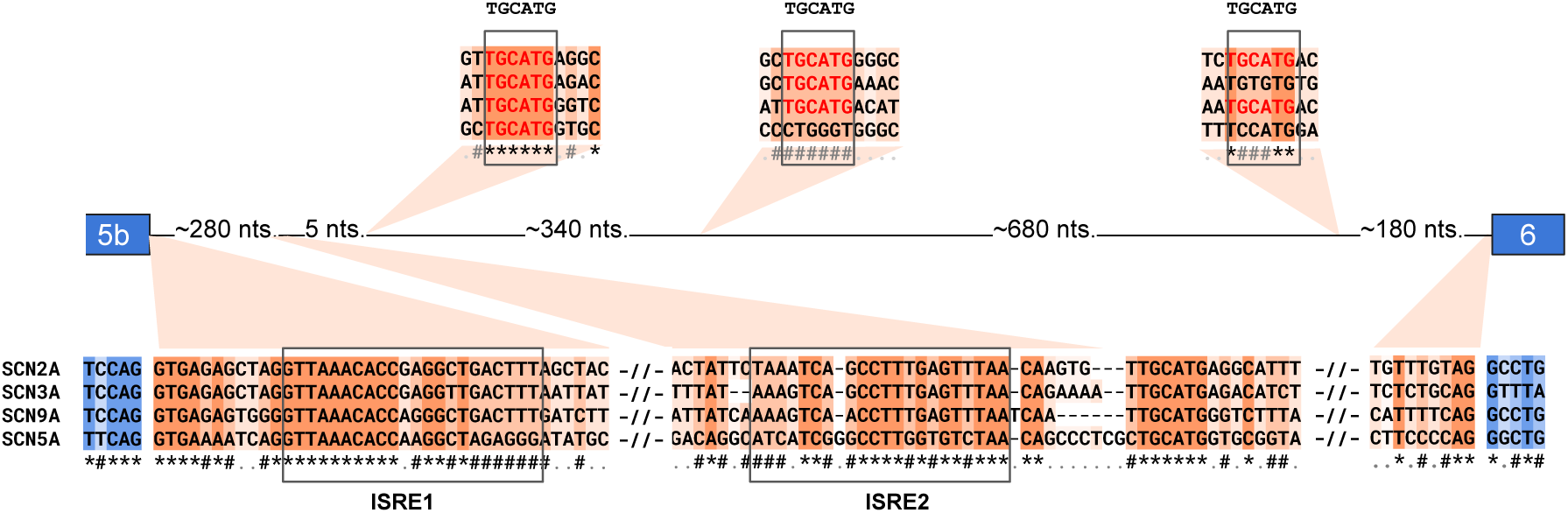
Multiple sequence alignment of intronic nucleotide sequences spanning between exons 5b and 6 in the human *SCN2A*, *SCN3A*, *SCN9A*, and *SCN5A* genes. A part of the alignment is shown (see Figures S4 and S5 for details). ‘*’ indicates columns with the same nucleotide in all sequences; # indicates columns with the same nucleotides in all but one sequence; ‘.’ indicates other columns. Insets above the gene line indicate consensus sequences (TGCATG) representing the canonical Rbfox2 binding motif.

**Figure 5:**
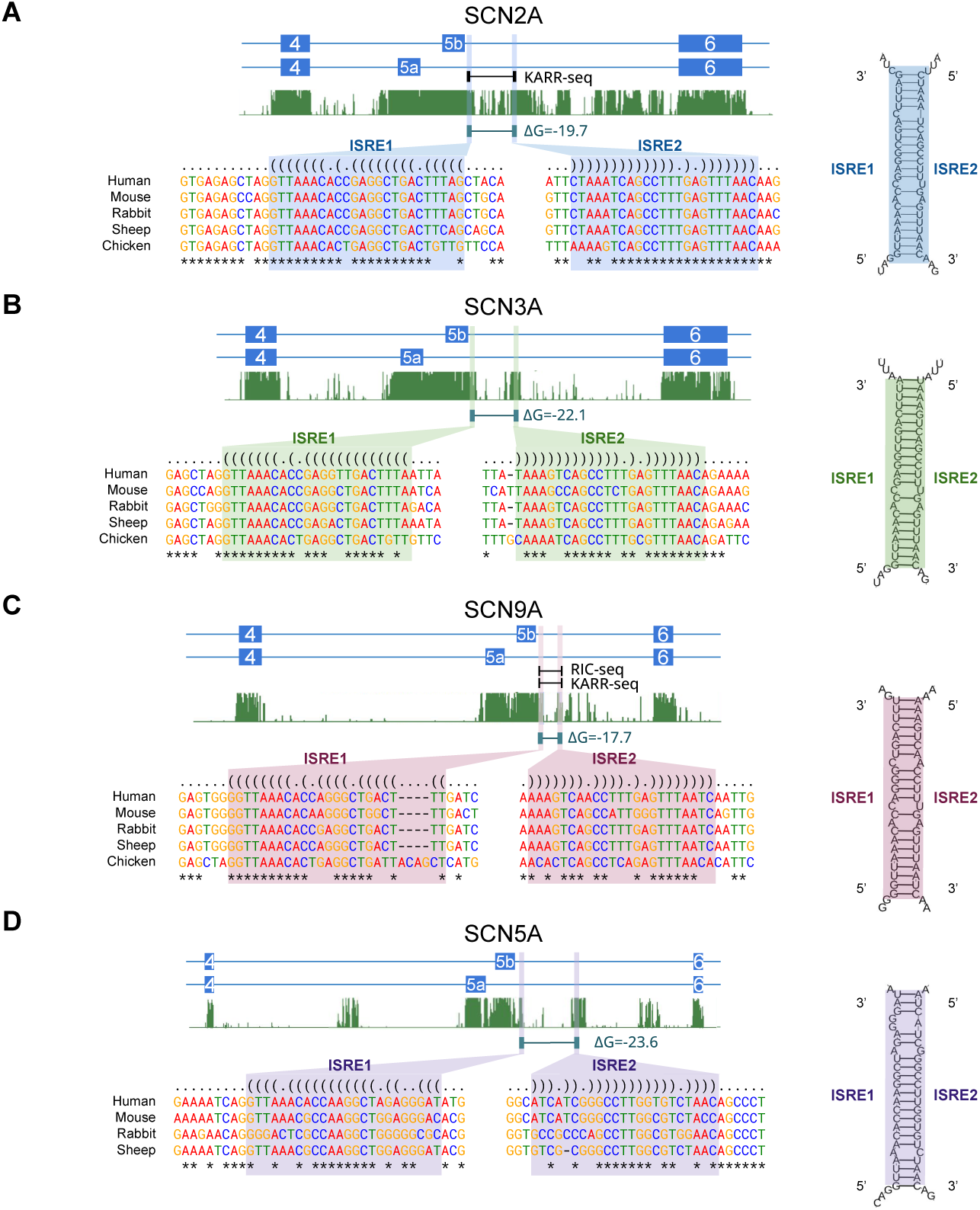
Intronic regulatory elements. **(A–D)** Left panels: genomic organization of exons 4-6 in the human *SCN2A*, *SCN3A*, *SCN9A*, and *SCN5A* genes, the locations of ISRE1 and ISRE2, their free energy of hybridization (Δ*G*), support by RIC-seq or KARR-seq (data pooled across cell lines, see Methods), 100 vertebrates conservation score by PhastCons (green track). Conserved nucleotides in multiple sequence alignments are indicated by asterisks. Right panels: RNA secondary structure formed by ISRE1 and ISRE2 binding.

Besides ISRE1 and ISRE2, the alignment highlighted three short homologous motifs with the consensus sequence TGCATG, one of which was located immediately downstream of ISRE2 (Figure 4). Considering that (U)GCAUG is the canonical binding motif of Rbfox proteins, we applied RBPNet, a deep learning model that predicts protein-RNA interactions by estimating CLIP-seq signal distribution from the RNA sequence [42], which confirmed sequence contexts of Rbfox2 binding in all four genes (Figure S6). These findings appear to be highly non-random since the probability of one particular hexamer occurring three or more times in an intron of length 1000 is below 0.5%.

We next checked whether ISRE1 and ISRE2 were listed in the genome-wide catalog of pairs of conserved complementary regions (PCCR) [43] or supported by proximity ligation assays targeting RNA structure (RIC-seq [44], PARIS [45], and KARR-seq [46]) in human cell lines (see Methods). Indeed, base pairings between ISRE1 and ISRE2 in *SCN2A*, *SCN3A*, *SCN5A*, and *SCN9A* were annotated as PCCRs with free energies (Δ*G*) ranging from -17 kcal/mol to -23.6 kcal/mol and confirmed by KARR-seq in *SCN2A*, and by both KARR-seq and RIC-seq in *SCN9A* (Figure 5A–D). The R-scape program [47] detected that ISRE1 and ISRE2 in the mammalian *SCN3A* and *SCN9A* genes acquired phylogenetically independent compensatory substitutions with the *E*-values of 0.45 and 0.06, respectively.

The base pairings between ISREs in *SCN1A* were listed as a PCCR, however with a weak hybridization energy (Δ*G* = −15 kcal/mol) due to nucleotide substitutions in ISRE1, while ISRE2 sequence was the same as in the other genes (Figure S7). Interestingly, the complementarity between ISRE1 and ISRE2 in chicken *SCN1A* was intact, but it was lost in sheep, indicating a separate evolutionary trajectory of ISRE1 in mammalian species. No sequences similar to ISRE1 or ISRE2 were detected in *SCN8A*, but there was an alternative complementary base pairing between intron and exon 5b sequences, which was supported by RIC-seq (Figure S7). Given that the divergence of *SCN8A* and *SCN1A*, *SCN2A*, *SCN3A*, *SCN9A* occurred later than that of *SCN8A* and *SCN5A*, the lack of common secondary structure may reflect its loss or evo-lutionary modification. Alignments to shuffled sequences showed that introns downstream of exon 5 in SCNα genes without exon 5 duplications contained no elements similar to ISRE1 and ISRE2 (Figure S8).

In sum, the identified sequences are common to introns of most SCNα genes with exon 5 duplications and capable of forming complementary base pairings. They are supported by RNA proximity ligation assays and, in some cases, by compensatory substitutions. As it appears highly unlikely to observe all this by pure chance, we chose to test experimentally whether ISRE1 and ISRE2 serve the purpose of forming RNA secondary structure.

### Validation of the ISRE impact on splicing

To test whether the interaction between ISRE influences MXE splicing, we quantitatively assessed the inclusion levels of exons 5a and 5b in human cell lines. According to Human Protein Atlas [48], *SCN9A* was the most expressed one among all human SCNα genes, and we chose to test it in the *Huh7* cells, where its expression level was the highest (Table S2). We designed antisense oligonucleotides (ASO) to disrupt ISRE1/ISRE2 base pairing and tested how they affect the inclusion levels of exons 5a and 5b in the endogenous *SCN9A* transcript. Since ASO reliably blocking ISRE1 could also interfere with the neighboring splice site, we focused on the ASO2 sequence that blocks ISRE2 (Figure 6A).

**Figure 6:**
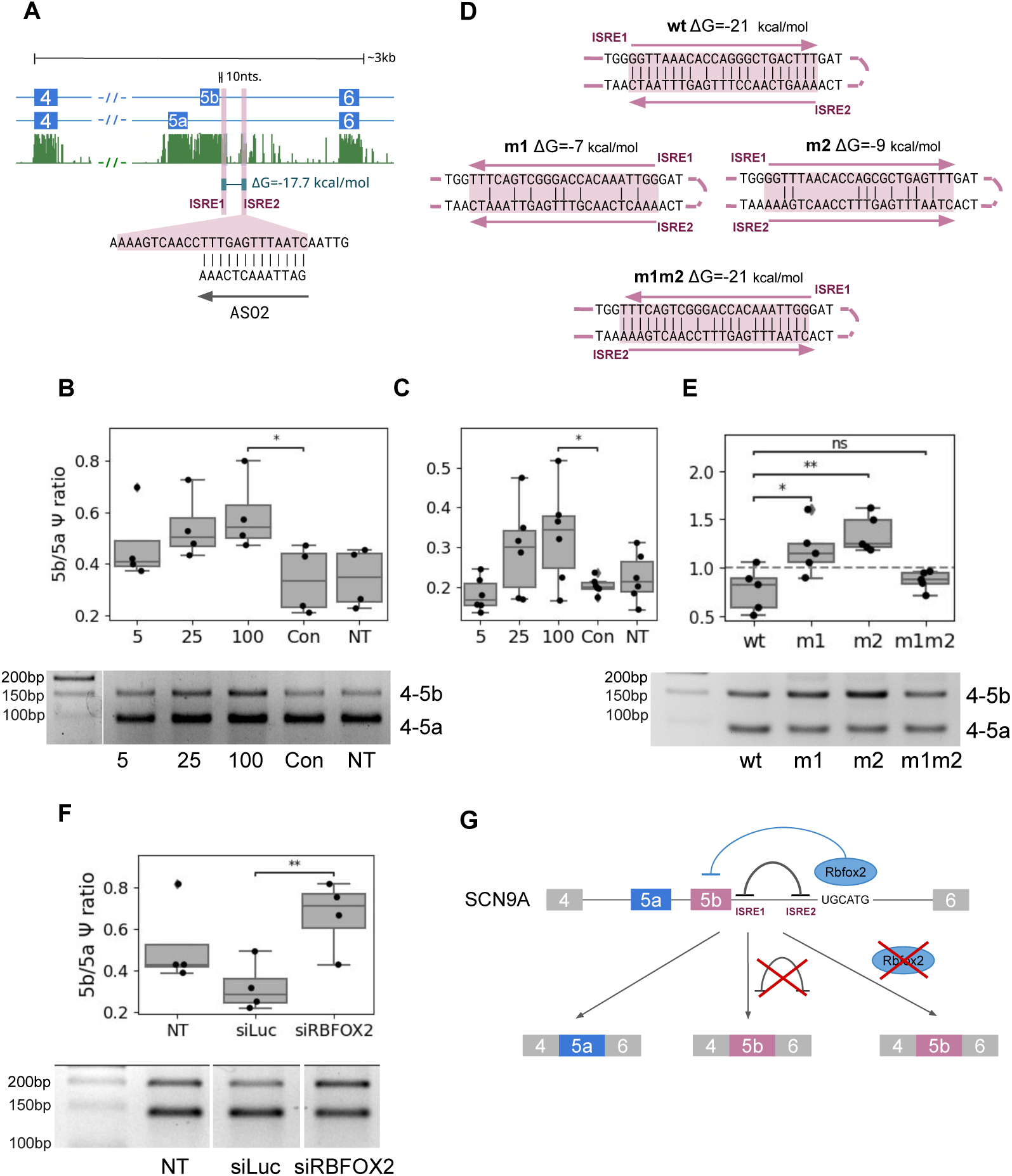
Experimental validation of RNA structure in the *SCN9A* gene and its impact on splicing. **(A)** The genomic organization of *SCN9A* exons 4–6, ISRE2 location, sequence and its corresponding ASO2. Δ*G* denotes the predicted base pairing energy; 10 nts is the distance between exon 5b and ISRE1. **(B)** RT-PCR quantification of exon 5b/5a Ψ ratio under ASO2 treatment in the *Huh7* cell line: gel image of RT-PCR products (bottom) and its densitometric quantitation (top). NT (non-treated control); Con (control ASO); 5, 25, 100 refer to ASO2 LNA concentration (nM). **(C)** RT-qPCR quantification of exon 5b/5a Ψ ratio under ASO2 treatment in the *Huh7* cell line; *x*-axis as in panel (B). **(D)** Disruptive and compensatory mutations (sequence reversal) in ISRE1 and ISRE2. wt — the wild type; m1, m2 — either ISRE1 or ISRE2 sequence is reversed as indicated by arrows; m1m2 — both ISRE1 and ISRE2 are reversed to restore base pairing. Δ*G* denotes the predicted base pairing energy. **(E)** RT-PCR quantification of exon 5b/5a Ψ ratio in single (m1 and m2) and double (m1m2) mutants. The *SCN9A* minigene was cloned in *HEK293T* cells (as in panel B). **(F)** RT-PCR quantification of exon 5b/5a Ψ ratio in siRNA-mediated KD of Rbfox2 (siRBFOX2) vs. KD of the luciferase gene (siLUC). **(G)** The proposed model of RNA bridge in Rbfox2-mediated regulation of exon 5b suppression. ‘*’ and ‘**’ denote statistically significant differences at the 5% and 1% significance levels, respectively (two-sample t-test); non-significant differences are labeled “ns”.

ASO2 was transfected into *Huh7* cells at three different concentration levels (5nM, 25nM, and 100nM) in comparison to the non-treated control and the treatment with a control ASO targeting an off-target site. Both qualitative assessment by endpoint RT-PCR (Figure 6B) and quantitative assessment of splicing by RT-qPCR (Figure 6C) showed that cells treated with ASO2 exhibit a significant concentration-dependent increase in the exon 5b/5a splicing ratio by approximately 15% and almost no variations in exon skipping and double exon inclusion (Figure S9). These results indicate that the ISRE2 sequence is important for maintaining the balance of splice isoforms with exons 5a and 5b without disrupting mutually exclusive splicing, resembling the pattern observed earlier for the *ATE1* gene [49].

Next, we created minigene constructs carrying a fragment of the *SCN9A* gene spanning between exons 4 and 6 with single mutations, in which the ISRE sequences were reversed to disrupt base pairing, and a double compensatory mutation, in which the base pairing was restored (Figure 6D). Transfection of these constructs into *HEK293T* cells, which lack endoge-nous *SCN9A* expression, revealed that disruptive mutations significantly increased the exon 5b/5a splicing ratio, while the double compensatory mutation reversed the splicing ratio to that of the wild type (Figure 6E). These results demonstrate that the balance of splice isoforms is, indeed, controlled by the complementary base pairing between ISRE1 and ISRE2.

Computational analysis of the potential protein binding sites within the intron between ex-ons 5b and 6 revealed three Rbfox2 binding sites, one of which was immediately proximal to ISRE2. The positioning of ISRE1 and ISRE2 suggests that RNA structure formed by their base pairing can function as an RNA bridge that brings distal *cis*-elements to the target exon [50]. Thus, we used siRNAs to perform a knockdown (KD) experiment in *Huh7* cells expressing endogenous *SCN9A* transcripts to check the impact of Rbfox2 inactivation on exon 5a/5b in-clusion. The exon 5b/5a splicing ratio significantly increased upon *RBFOX2* KD, indicating that both the ISRE1/ISRE2 interaction and Rbfox2 binding are important in suppressing the inclusion of exon 5b (Figure 6F).

### Evolutionary origin of ISRE in SCN**α** genes

The remarkable similarity of ISRE sequences in the human SCNα genes with exon 5 duplica-tions suggests that these regulatory elements may have existed in their common ancestor. To test this hypothesis, we aligned intronic sequences downstream of exon 5b in all genes with anno-tated or predicted exon 5 duplications from human, mouse, sheep, chicken, frog, zebrafish, ele-phant shark, lancelet, and *Ciona*. In most species, the homologs of the human *SCN1A*, *SCN2A*, *SCN3A*, *SCN9A*, and *SCN5A* genes maintained ISRE sequences, with the highest degree of con-servation observed in amniotes (Figure 7A). The organization of these loci completely matched that of the human ones with the exception of chicken *SCN5A* and sheep *SCN1A*, both of which lost ISRE2, and mouse *SCN1A*, which is lacking both exon duplication and ISREs.

**Figure 7:**
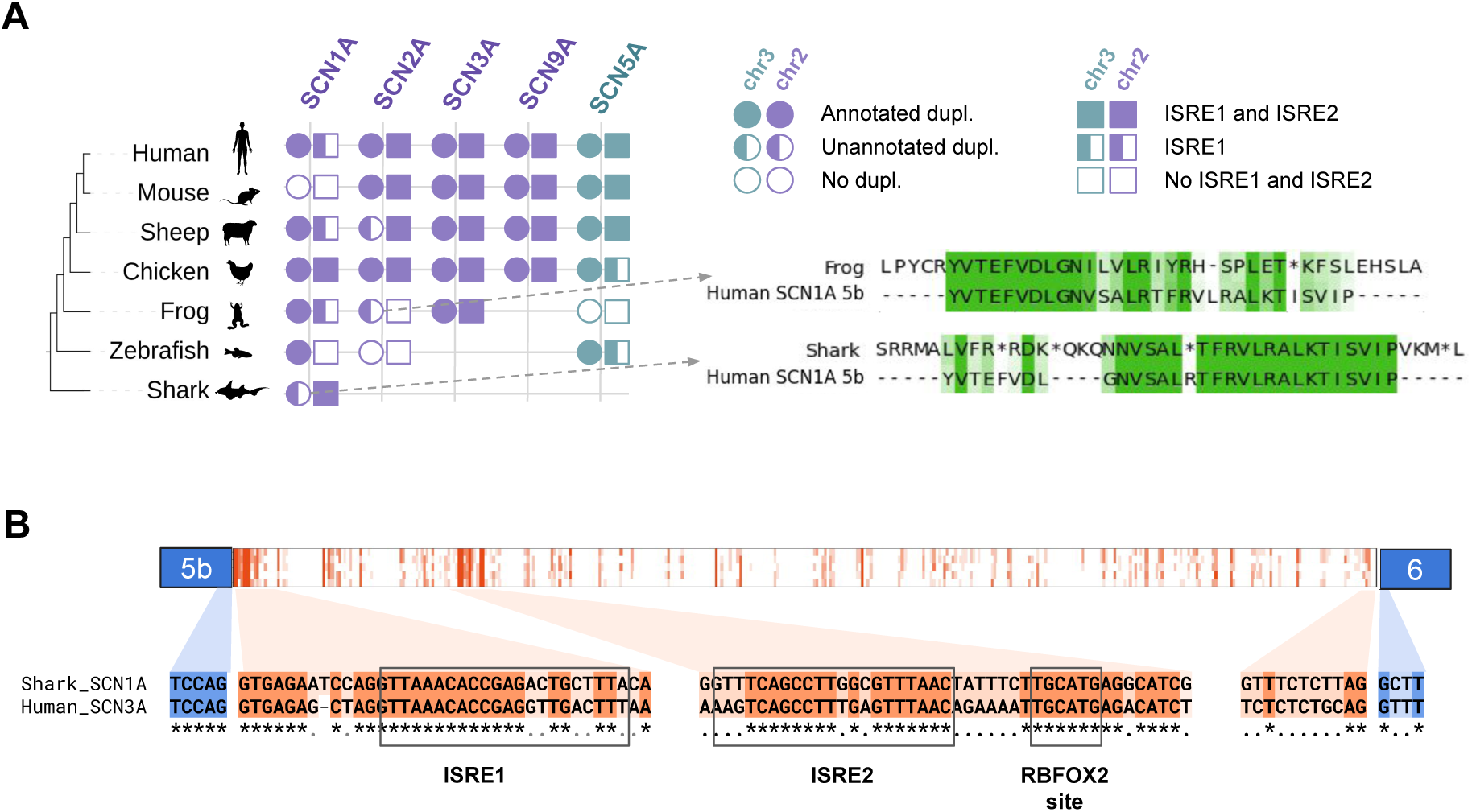
Common RNA structures in vertebrate SCNα genes. **(A)** SCNα genes, their exon 5 duplications and presence of ISRE1/ISRE2 sequences. **(B)** Top: a heatmap representation of the alignment of the *SCN1A* intron between exons 5b and 6 in elephant shark with the *SCN3A* introns between exons 5b and 6 in human, mouse, chicken and frog. Shades of red color indicate similarity between the aligned positions. Bottom: the alignment of the elephant shark *SCN1A* and human *SCN3A* indicates the ancient origin of ISRE1, ISRE2, and the Rbfox2 binding site. ‘*’ and ‘.’ are as in Figure 4.

Notable patterns emerged in even more distant taxa. For example, in frogs, the gene cluster containing *SCN1A*, *SCN2A*, and *SCN3A* maintained exon 5 duplication and *SCN3A* retained complete ISRE sequences, while the *SCN5A* gene lacked both exon 5 duplication and ISREs. In zebrafish, *SCN5A* showed an exon 5 duplication pattern homologous to that in humans but lacked ISRE2. Among zebrafish *SCN1A* homologs, one maintained exon 5 duplication but lacked both ISREs, while another had neither duplication nor conserved intronic regions. An interesting case was found in elephant shark (*C. milii*), where *SCN1A* homolog had both exon 5 duplication and ISRE sequences, while other SCNα gene homologs in sharks lacked these features (Figure 7B). The shark *SCN1A* homolog was phylogenetically the closest among all shark *SCN*α genes to the lineage bearing both exon 5 duplication and ISREs. Interestingly, the number of base pairs between ISRE1 and ISRE2 in the shark *SCN1A* and in the human *SCN3A* were the same, but their positions were not.

These findings suggest that the elephant shark represents, perhaps, the most distant from the human living organism possessing both exon 5b duplication and ISRE sequences capable of forming RNA structure, which potentially regulate alternative splicing of exons 5b/5a. These structural elements may have been present in the common ancestor of all vertebrates with jaws, whereas exon duplications appeared earlier in the common ancestor of all chordates. This evolutionary dynamics highlights the important role of complementary intronic sequences in regulating SCNα gene splicing throughout vertebrate history.

## Discussion

Current evidence suggests that MXE splicing is crucial for shaping proteomic diversity and controlling transcript isoform expression in a tissue- and developmental stage-specific manner [51, 52]. It is a strictly coordinated mechanism, in which one and only one of two or more consecutive exons is included in the mature mRNA. In some cases, the mutually exclusive splicing pattern is characteristic to entire paralogous gene families, while in others each gene specifically conserves just one particular exon [53]. RNA structure is often associated with MXE regulation [54], but typically such regulatory mechanism is studied in just one paralog, e.g., in the *TPM2* gene from the tropomyosin gene family [55]. It is therefore natural to ask whether a group of paralogs possess regulatory elements that were present in the ancestral gene and maintained by evolution after gene duplication. Here, we present a comprehensive analysis of splicing regulation in the SCNα gene family, shedding light on splicing evolution and providing a framework for identifying conserved regulatory elements through paralog comparison.

The intronic splicing regulatory elements identified in this study are homologous across paralogs and exhibit multiple characteristic features of functional RNA structures such as compensatory substitutions and support by RNA proximity ligation assays. They strongly resemble splicing regulatory elements in FGFR1 and FGFR2 genes, which also display evolutionarily conserved, homologous sequence patterns [23]. In the *SCN9A* gene, single mutations disrupting the base pairing between ISRE1 and ISRE2 promote exon 5b inclusion, while double compensatory mutation restores the original splicing ratio. Furthermore, the disruption of ISRE2 accessibility by ASO altered the 5b/5a isoform ratio in the endogenous *SCN9A* transcript without the loss of mutual exclusivity, recapitulating observations in the *ATE1* gene, where a long-range RNA structure similarly regulates the splice isoform balance [49]. These findings support the hypothesis that ISRE1 and ISRE2 represent an evolutionarily ancient and functionally important mechanism for controlling the ratio of splice isoforms in the SCNα gene family through RNA structure-mediated regulation.

However, several factors challenge a straightforward interpretation of the ISRE1 and ISRE2 forming RNA structure in other members of the SCNα gene family. While these regulatory elements are complementary in most SCNα paralogs with exon 5 duplication, their complementarity in SCN1A is notably weaker due to sequence divergence of ISRE1, a puzzling observation given a high level of sequence conservation in ISRE2. Such structural flexibility still may reflect adaptation to tissue-specific regulatory requirements. Unlike other known RNA structures with impact on splicing [56], the predicted ISRE1/ISRE2 interaction lacks extended continuous helices, instead forming shorter, more diffuse interactions. This leaves a possibility for other regulatory actors such as RNA-binding proteins.

The Rbfox proteins are widely known to be master regulators of alternative splicing in many tissues including skeletal muscle, heart, and brain [57, 58, 59, 60, 61, 62]. In voltage-gated sodium channel genes, their impact on alternative splicing of duplicated exons was studied mostly in *SCN8A* and *SCN5A*, where exons 5a and 5b are referred to as ‘N’ (neonatal) and ‘A’ (adult), respectively. Surprisingly, the Rbfox proteins act differently in paralogs: Rbfox2 overexpression suppresses the inclusion of exon 5b in cardiac *SCN5A* [63], while the inclusion of 5b in *SCN8A* is suppressed not by overexpression but by the loss of Rbfox2 or Rbfox1, which has the same RNA-binding motif as Rbfox2 [59, 58]. Thus, two opposite perturbations of Rbfox expression led to the same splicing outcome in different paralogs. Furthermore, different splice isoforms of Rbfox2 differently impact the choice of 5a and 5b in *SCN5A* [63].

Our findings suggest a mechanistic model, in which the RNA structure formed by ISRE1 and ISRE2 acts as an RNA bridge, a structural element that brings the RBP binding site closer to the regulated exon (Figure 6G). For instance, an RNA bridge is necessary for Rbfox-mediated inclusion of exon 11a in the ENAH gene [50]. As we have shown for *SCN9A*, both disruption of ISRE1/ISRE2 base pairing and Rbfox2 depletion promote exon 5b inclusion, which implies that both RNA structure and Rbfox2 binding are needed to suppress the isoform with exon 5b. The difference between ENAH exon 11a and *SCN9A* exon 5b is that, in the latter case, Rbfox2 acts not as an activator but as a suppressor of splicing. Such opposite functions are not uncommon among RBPs, e.g., PTBP1 may act as both a splicing repressor and activator depending on the number of binding sites and their locations relative to the target exon [64, 65].

Yet, the mechanism underlying mutual exclusive choice of exons 5a and 5b in SCNα par-alogs with exon 5 duplication remains unresolved. Our findings suggest that it is not the NMD pathway that removes splice isoforms in which exons 5a and 5b are both included or skipped. If it were NMD, we should have seen their upregulation in NMD inactivation experiments and a comparable proportion of skipped and 5a5b isoforms in other conditions, but the 5a5b isoform was almost never observed (Figure 3). Although a NMD-dependent degradation was proposed to regulate mutual exclusive splicing in pairs of MXEs that are not multiples of three in length [13], in the case of SCNα genes it must be a different mechanism, possibly another group of RNA structure elements that was not detected by comparative sequence analysis.

Taken together, our findings not only highlight the common mechanism of splicing regulation in the SCNα gene family, but they also demonstrate the power of comparative genomics in deciphering splicing regulatory mechanisms in paralogs. In most SCNα genes (with the exception of *SCN9A*), the complementarity between ISRE1 and ISRE2 cannot be found using single-sequence RNA folding tools such as MFOLD [66], while phylogenetic methods don’t have enough statistical power due to extreme level of sequence conservation and absence of not only compensatory, but any substitutions at all. In contrast, paralogy-based approaches can underscore functional base-paired regions by contrasting sequence divergence with preservation of regulatory elements. Future studies applying this strategy to other gene families may reveal many more similarly hidden regulatory architectures.

## Methods

### Gene sequences and search for unannotated exon duplications

The nucleotide sequences and exon-intron boundaries of annotated transcript isoforms of homologous SCNα genes across 13 different species were obtained from the Ensembl and NCBI databases (Table S3). To identify unannotated SCNα genes in other organisms, tblastn (from BLAST v2.12.0+ [67]) alignments were performed against the respective genomes with the an-notated SCNα gene aminoacid sequence as a query. The MAFFT v7.490 multiple alignment tool [68] was used to confirm the homology of the selected genes. Consecutive exons of the same gene with high aminoacid sequence identity (>30%) were classified as tandemly duplicated exons. In order to identify unannotated exon duplications, the translated sequence of each human exon (5a or 5b) was aligned against the intronic sequences flanking its homolog in the target species in all possible reading frames using tblastn tool. Additionally, we aligned exons in the target species against their own flanking introns. To assess the significance of the predicted exon homology, we used the false positive rate (FPR) metric computed from realignments of the query sequence to 1000 random permutations of the target aminoacid sequence by counting the number of times when the random permutation had a higher alignment score than the actual alignment. Only exons with FPR below 5% are reported throughout this study.

### The construction of phylogenetic trees

Multiple sequence alignments were constructed separately for two groups of amino acid se-quences, the major protein isoform of each SCNα gene and for the translated sequences of an-notated and newly discovered exons 5a and 5b using Muscle5 program with the default settings [69]. The aminoacid sequence of the major protein isoform was used to construct a phylogenetic tree. The optimal evolutionary model was selected using IQ-TREE v3.0.1 ModelFinder, with the JTT+R8 model identified as most appropriate [70]. Phylogenetic trees were constructed in IQ-TREE using the selected evolutionary model. Branch support values were assessed through ultrafast bootstrap analysis with 1000 replicates.

### Identification and analysis of ISRE

To identify ISRE, we performed multiple sequence alignment of homologous introns flanking mutually exclusive exons. Blocks of homologous sequences of at least 10 nts in length were extracted from these alignments and subsequently analyzed using PrePH [43] and RNAup v2.3.5 program from the Vienna RNA package [71] programs with default settings to predict complementary base pairings. For compensatory mutation analysis, we applied the R-scape v2.0.0 software [47] to blocks of multiple sequence alignments of 30 mammalian species after merging through a 10-nt spacer as explained earlier [43]. The E-value of the structure (*E*) was calculated as the product of E-values of its constituent base pairs that were provided by R-scape.

### RNA-seq data and percent-spliced-in (PSI) metric

The poly(A)+ RNA-seq data of human tissues (Table S4) from the GTEx project [72] and RNA-seq data of matched cancer and normal human tissues (Table S5) from The Cancer Genome Atlas (TCGA) consortium [73] were obtained and processed as explained earlier [65, 74]. The poly(A)+ RNA-seq data from the study of mammalian organ development were obtained from [37] in BAM format (Table S6). The poly(A)+ RNA-seq data from the study of mouse embryonic transcriptomes [38] were downloaded from the ENCODE Consortium portal in BAM format. The poly(A)+ RNA-seq data from i3Neurons with NMD inhibition via single and double knockdowns of XRN1, SMG6, and UPF1 [75] and from A549 cells with NMD inhibition by cycloheximide [74] were downloaded from the Gene Expression Omnibus website in FASTQ format under the accession numbers GSE307054 and GSE270310, respectively, and aligned to the GRCh38 human genome assembly using STAR aligner v2.7.10a with default parameters [76]. Short read alignments were subsequently analyzed by the IPSA pipeline with the default settings [77]. The obtained split read counts were used to compute the Percent-Spliced-In (PSI, Ψ) metric, defined as the number of split reads supporting exon inclusion as a fraction of the collective number of split reads supporting exon inclusion and skipping.

### RNA proximity ligation data

The following RNA proximity ligation data were used: RNA *in situ* conformation sequenc-ing (RIC-seq) from the *HeLa*, *K562*, *HepG2*, *IMR90*, *hNPC*, *H1*, *GM12878* cells as described [78]; kethoxal-assisted RNA–RNA interaction sequencing (KARR-seq) in the *F123*, *HEK293T*, *HepG2*, *K562*, *A549* cells [46]; and psoralen analysis of RNA interactions and structures (PARIS) in the *HEK293*, *HeLa*, and *SH-SY5Y* cells. All RNA contacts from each experiment type were polled and represented via UCSC Genome Browser track as explained in [79].

### Cell lines

The *Huh7* cell line was used to determine the expression of splice isoforms with qPCR. The cell line was cultured in DMEM/F-12 (Paneco LLC, Russia) medium supplemented with 10% fetal bovine serum (INTL Kang, China), penicillin (50 IU/ml), and streptomycin (0.05 mg/ml) (Thermo Fisher Scientific, USA) at 37°C and 5% CO_2_ in the atmosphere.

### Minigene construction and mutagenesis

Whole genomic DNA of the *HEK293T* cell line was isolated using LumiSpin® UNI, DNA Iso-lation Spin Kit for Any Sample (Lumiprobe, Russia). A fragment of the *SCN9A* gene spanning the interval from exon 4 to exon 6 was amplified from the genomic DNA using Q5 High-Fidelity DNA Polymerase (New England Biolabs) and inserted downstream of the cytomegalovirus (CMV) promoter of the pRK5 vector (kindly provided by Prof. P.M. Rubtsov). The minigene was made using the CloneExpress Ultra One Step cloning kit (Vazyme, China). Mutations in all minigenes were introduced by polymerase chain reaction (PCR) using phosphorylated primers (Table S7) carrying desired mutations with subsequent ligation by T4 DNA ligase (Evrogen, Russia) using Rapid Ligation protocol. All constructs were confirmed by sequencing and re-striction analysis.

### RBFOX2 inactivation experiments

*Huh7* cells were plated at a density of 70,000 cells per well on a 24-well plate. The transfection of siRNA against *RBFOX2* into cells was performed using Lipofectamine RNAiMAX (Invit-rogen) in OptiMEM serum-reduced media (Gibco) at siRNA concentration of 10nM. The cells transfected with non-targeting luciferase siRNA were used as a negative control. The siRNA se-quences are listed in Table S8. Cells were harvested after 48h of treatment for further analysis. The knockdown efficiency was ¿90% as confirmed by RT-qPCR (primers in Table S9).

### Antisense oligonucleotides (ASO)

LNA/DNA mixmers (Syntol LLC, Russia) were used as antisense oligonucleotides to block complementary base pairs within RNA. ASO had a phosphorothioated backbone to protect against cellular nucleases. The sequence of ASO2 and the control sequence were +G*A*+T*T* +A*A*+A*C*+T*C*+A*A*+A and +T*G*+G*A*+A*G*+T*C*+T*T*+C*G*+T, respec-tively. In these sequences, G, A, T, and C denote DNA bases; +A and +G denote LNA bases; G*, A*, T*, and C* denote phosphorothioated DNA bases; +G*, +A*, +T*, and +C* denote phosphorothioated LNA bases.

### ASO transfection

Approximately 200,000 *Huh7* cells were seeded into a 12-well plate and cultured for 24 hours. ASO transfection into the cells was performed using Lipofectamine RNAiMAX (Invitrogen), at ASO concentrations of 5, 25, and 100 nM. 3 µl of Lipofectamine and the appropriate concentration of LNA/DNA oligonucleotide were used per well, the lipid-RNA complex was dissolved in Opti-MEM Reduced Serum Medium (Gibco) and added to the cells according to the manufacturer’s protocol. The cells were then incubated for 24 hours before harvesting.

### RNA isolation

Total RNA was isolated using RUplus Column total RNA isolation kit (Biolabmix, Russia). One microgram of total RNA was first subjected to RNase-free DNase I digestion (Thermo Fisher Scientific, USA) at 37°C for 30 min to remove contaminating genomic DNA. Next, 500 ng of total RNA was used for complementary DNA (cDNA) synthesis using Maxima First Strand cDNA Synthesis Kit for RT-qPCR (Thermo Fisher Scientific, USA) to a final volume of 5 µl. cDNA was diluted 1:10 with nuclease-free water for quantitative PCR (qPCR) and reverse transcription PCR (RT-PCR) analysis.

### RT-qPCR

RT-qPCR reactions were performed in triplicates in a final volume of 12 µl in 96-well plates with 420 nM gene-specific primers and 2 µl of cDNA using 5X qPCRmix-HS SYBR reaction mix (Evrogen, Russia). Primers for RT-qPCR are listed in Table S9. Since exons 5a and 5b are both 92-nts-long, the primer positions were chosen to yield PCR products of different size (Figure S10). A sample without reverse transcriptase enzyme was included as control to verify the absence of genomic DNA contamination. Amplification of the targets was carried out on CFX96 Real-Time System (Bio-Rad, USA), with following parameters: 95°C for 2 min, fol-lowed by 42 cycles at 95°C for 30 s, 57°C for 30 s and 72°C for 30 s, ending at 72°C for 5 min. Gene and gene isoform expression change was calculated using an estimate of the amplification efficiency value.

### Statistical tests

The data were analyzed using python version 3.8.2 and R statistics software version 3.6.3. Two-tailed t-test (with Bonferroni correction in the case of multiple comparisons) was performed if no departure from population normality was observed. Non-parametric tests were performed using normal approximation with continuity correction. In all figures, the significance levels 5% and 0.1% are denoted by * and ***, respectively.

## Data and code availability

All scripts, command line calls, and processed data supporting this study are publicly available through Zenodo repository [80].

## Authors’ contributions

EC and DP designed the study; EC performed data analysis; MV and DS performed the experi-ments; EC and DP wrote the first draft of the manuscript. All authors edited the final version of the manuscript.

## Acknowledgments

The authors thank Prof. O.A. Dontsova for insightful discussions.

## References

[1] Wang, J., Ou, S.-W., and Wang, Y.-J. (2017) Distribution and function of voltage-gated sodium channels in the nervous system. Channels (Austin), 11, 534–554.

[2] Stuhmer, W., Conti, F., Suzuki, H., Wang, X. D., Noda, M., Yahagi, N., Kubo, H., and Numa, S. (1989) Structural parts involved in activation and inactivation of the sodium channel. Nature, 339, 597–603.

[3] Catterall, W. A., Goldin, A. L., and Waxman, S. G. (2005) International Union of Phar-macology. XLVII. Nomenclature and structure-function relationships of voltage-gated sodium channels. Pharmacol Rev, 57, 397–409.

[4] Alabi, A. A., Bahamonde, M. I., Jung, H. J., Kim, J. I., and Swartz, K. J. (2007) Portability of paddle motif function and pharmacology in voltage sensors. Nature, 450, 370–5.

[5] Mantegazza, M., Cestele, S., and Catterall, W. A. (2021) Sodium channelopathies of skele-tal muscle and brain. Physiol Rev, 101, 1633–1689.

[6] Heighway, J., Sedo, A., Garg, A., Eldershaw, L., Perreau, V., Berecki, G., Reid, C. A., Petrou, S., and Maljevic, S. (2022) Sodium channel expression and transcript variation in the developing brain of human, Rhesus monkey, and mouse. Neurobiol Dis, 164, 105622.

[7] Liang, L., Fazel Darbandi, S., Pochareddy, S., Gulden, F. O., Gilson, M. C., Sheppard, B. K., Sahagun, A., An, J.-Y., Werling, D. M., Rubenstein, J. L. R., Sestan, N., Bender, K. J., and Sanders, S. J. (2021) Developmental dynamics of voltage-gated sodium channel isoform expression in the human and mouse brain. Genome Med, 13, 135.

[8] Pang, P. D., Alsina, K. M., Cao, S., Koushik, A. B., Wehrens, X. H. T., and Cooper, T. A. (2018) CRISPR-Mediated Expression of the Fetal Scn5a Isoform in Adult Mice Causes Conduction Defects and Arrhythmias. J Am Heart Assoc, 7, e010393.

[9] Raymond, C. K., Castle, J., Garrett-Engele, P., Armour, C. D., Kan, Z., Tsinoremas, N., and Johnson, J. M. (2004) Expression of alternatively spliced sodium channel alpha-subunit genes. Unique splicing patterns are observed in dorsal root ganglia. J Biol Chem, 279, 46234–41.

[10] Freyermuth, F., Rau, F., Kokunai, Y., Linke, T., Sellier, C., Nakamori, M., Kino, Y., Arandel, L., Jollet, A., Thibault, C., Philipps, M., Vicaire, S., Jost, B., Udd, B., Day, J. W., Duboc, D., Wahbi, K., Matsumura, T., Fujimura, H., Mochizuki, H., Deryckere, F., Kimura, T., Nukina, N., Ishiura, S., Lacroix, V., Campan-Fournier, A., Navratil, V., Chautard, E., Auboeuf, D., Horie, M., Imoto, K., Lee, K.-Y., Swanson, M. S., de Munain, A. L., Inada, S., Itoh, H., Nakazawa, K., Ashihara, T., Wang, E., Zimmer, T., Furling, D., Takahashi, M. P., and Charlet-Berguerand, N. (2016) Splicing misregulation of SCN5A contributes to cardiac-conduction delay and heart arrhythmia in myotonic dystrophy. Nat Commun, 7, 11067.

[11] Menezes, L. F. S., Sabiá Júnior, E. F., Tibery, D. V., Carneiro, L. D. A., and Schwartz, E. F. (2020) Epilepsy-Related Voltage-Gated Sodium Channelopathies: A Review. Front Pharmacol, 11, 1276.

[12] Munyao, W., Rahman, M. M., Sabzanov, S. A., Chu, E. H., Wang, R., Wang, Z., Yu, Y., and Ruggiu, M. (2025) Alternative Splicing and CaV-Associated Channelopathies. Wiley Interdiscip Rev RNA, 16, e70016.

[13] Jin, Y., Dong, H., Shi, Y., and Bian, L. (2018) Mutually exclusive alternative splicing of pre-mRNAs. Wiley Interdiscip Rev RNA, 9, e1468.

[14] May, G. E., Olson, S., McManus, C. J., and Graveley, B. R. (2011) Competing RNA secondary structures are required for mutually exclusive splicing of the Dscam exon 6 cluster. RNA, 17, 222–9.

[15] Wang, X., Li, G., Yang, Y., Wang, W., Zhang, W., Pan, H., Zhang, P., Yue, Y., Lin, H., Liu, B., Bi, J., Shi, F., Mao, J., Meng, Y., Zhan, L., and Jin, Y. (2012) An RNA architectural locus control region involved in Dscam mutually exclusive splicing. Nat Commun, 3, 1255.

[16] Hong, W., Shi, Y., Xu, B., and Jin, Y. (2020) RNA secondary structures in Dscam1 mutually exclusive splicing: unique evolutionary signature from the midge. RNA, 26, 1086–1093.

[17] Hatje, K. and Kollmar, M. (2013) Expansion of the mutually exclusive spliced exome in Drosophila. Nat Commun, 4, 2460.

[18] Ivanov, T. M. and Pervouchine, D. D. (2018) An Evolutionary Mechanism for the Generation of Competing RNA Structures Associated with Mutually Exclusive Exons. Genes (Basel), 9, 356.

[19] Ivanov, T. M. and Pervouchine, D. D. (2022) Tandem Exon Duplications Expanding the Alternative Splicing Repertoire. Acta Naturae, 14, 73–81.

[20] Parker, B. J., Moltke, I., Roth, A., Washietl, S., Wen, J., Kellis, M., Breaker, R., and Pedersen, J. S. (2011) New families of human regulatory RNA structures identified by comparative analysis of vertebrate genomes. Genome Res, 21, 1929–43.

[21] Soll, D. R. and Waddell, D. R. (1975) Morphogenesis in the slime mold Dictyostelium discoideum. 1. The accumulation and erasure of “morphogenetic information”. Dev Biol, 47, 292–302.

[22] Mistry, N., Harrington, W., Lasda, E., Wagner, E. J., and Garcia-Blanco, M. A. (2003) Of urchins and men: evolution of an alternative splicing unit in fibroblast growth factor receptor genes. RNA, 9, 209–17.

[23] Muh, S. J., Hovhannisyan, R. H., and Carstens, R. P. (2002) A Non-sequence-specific double-stranded RNA structural element regulates splicing of two mutually exclusive exons of fibroblast growth factor receptor 2 (FGFR2). J Biol Chem, 277, 50143–54.

[24] Petrova, M., Margasyuk, S., Vorobeva, M., Skvortsov, D., Dontsova, O. A., and Pervouchine, D. D. (2024) BRD2 and BRD3 genes independently evolved RNA structures to control unproductive splicing. NAR Genom Bioinform, 6, lqad113.

[25] Xu, L., Ding, X., Wang, T., Mou, S., Sun, H., and Hou, T. (2019) Voltage-gated sodium channels: structures, functions, and molecular modeling. Drug Discov Today, 24, 1389–1397.

[26] Gazina, E. V., Richards, K. L., Mokhtar, M. B. C., Thomas, E. A., Reid, C. A., and Petrou, S. (2010) Differential expression of exon 5 splice variants of sodium channel alpha subunit mRNAs in the developing mouse brain. Neuroscience, 166, 195–200.

[27] de Lera Ruiz, M. and Kraus, R. L. (2015) Voltage-Gated Sodium Channels: Structure, Function, Pharmacology, and Clinical Indications. J Med Chem, 58, 7093–118.

[28] Catterall, W. A. (2014) Structure and function of voltage-gated sodium channels at atomic resolution. Exp Physiol, 99, 35–51.

[29] Zakon, H. H. (2012) Adaptive evolution of voltage-gated sodium channels: the first 800 million years. Proc Natl Acad Sci U S A, 109 **Suppl 1**, 10619–25.

[30] Liavas, A., Lignani, G., and Schorge, S. (2017) Conservation of alternative splicing in sodium channels reveals evolutionary focus on release from inactivation and structural insights into gating. J Physiol, 595, 5671–5685.

[31] Sanchez-Sandoval, A. L., Hernández-Plata, E., and Gomora, J. C. (2023) Voltage-gated sodium channels: from roles and mechanisms in the metastatic cell behavior to clinical potential as therapeutic targets. Front Pharmacol, 14, 1206136.

[32] Saavedra-Rodriguez, K., Maloof, F. V., Campbell, C. L., Garcia-Rejon, J., Lenhart, A., Penilla, P., Rodriguez, A., Sandoval, A. A., Flores, A. E., Ponce, G., Lozano, S., and Black, W. C. (2018) Parallel evolution of vgsc mutations at domains IS6, IIS6 and IIIS6 in pyrethroid resistant Aedes aegypti from Mexico. Sci Rep, 8, 6747.

[33] Yuan, H., Shan, W., Zhang, Y., Yan, H., Li, Y., Zhou, Q., Dong, H., Tao, F., Liu, H., Leng, P., Peng, H., and Ma, Y. (2023) High frequency of Voltage-gated sodium channel (VGSC) gene mutations in Aedes albopictus (Diptera: Culicidae) suggest rapid insecticide resistance evolution in Shanghai, China. PLoS Negl Trop Dis, 17, e0011399.

[34] Yeo, G. W., Van Nostrand, E. L., and Liang, T. Y. (2007) Discovery and analysis of evolutionarily conserved intronic splicing regulatory elements. PLoS Genet, 3, e85.

[35] Altschul, S. F., Gish, W., Miller, W., Myers, E. W., and Lipman, D. J. (1990) Basic local alignment search tool. J Mol Biol, 215, 403–10.

[36] Copley, R. R. (2004) Evolutionary convergence of alternative splicing in ion channels. Trends Genet, 20, 171–6.

[37] Mazin, P. V., Khaitovich, P., Cardoso-Moreira, M., and Kaessmann, H. (2021) Alternative splicing during mammalian organ development. Nat Genet, 53, 925–934.

[38] He, P., Williams, B. A., Trout, D., Marinov, G. K., Amrhein, H., Berghella, L., Goh, S.-T., Plajzer-Frick, I., Afzal, V., Pennacchio, L. A., Dickel, D. E., Visel, A., Ren, B., Hardison, R. C., Zhang, Y., and Wold, B. J. (2020) The changing mouse embryo transcriptome at whole tissue and single-cell resolution. Nature, 583, 760–767.

[39] Bahcheli, A. T., Min, H.-K., Bayati, M., Zhao, H., Fortuna, A., Dong, W., Dzneladze, I., Chan, J., Chen, X., Guevara-Hoyer, K., Dirks, P. B., Huang, X., and Reimand, J. (2024) Pan-cancer ion transport signature reveals functional regulators of glioblastoma aggression. EMBO J, 43, 196–224.

[40] House, C. D., Vaske, C. J., Schwartz, A. M., Obias, V., Frank, B., Luu, T., Sarvazyan, N., Irby, R., Strausberg, R. L., Hales, T. G., Stuart, J. M., and Lee, N. H. (2010) Voltage-gated Na+ channel SCN5A is a key regulator of a gene transcriptional network that controls colon cancer invasion. Cancer Res, 70, 6957–67.

[41] Igci, Y. Z., Bozgeyik, E., Borazan, E., Pala, E., Suner, A., Ulasli, M., Gurses, S. A., Yum-rutas, O., Balik, A. A., and Igci, M. (2015) Expression profiling of SCN8A and NDUFC2 genes in colorectal carcinoma. Exp Oncol, 37, 77–80.

[42] Horlacher, M., Wagner, N., Moyon, L., Kuret, K., Goedert, N., Salvatore, M., Ule, J., Gagneur, J., Winther, O., and Marsico, A. (2023) Towards in silico CLIP-seq: predicting protein-RNA interaction via sequence-to-signal learning. Genome Biol, 24, 180.

[43] Kalmykova, S., Kalinina, M., Denisov, S., Mironov, A., Skvortsov, D., Guigo, R., and Per-vouchine, D. (2021) Conserved long-range base pairings are associated with pre-mRNA processing of human genes. Nat Commun, 12, 2300.

[44] Cai, Z., Cao, C., Ji, L., Ye, R., Wang, D., Xia, C., Wang, S., Du, Z., Hu, N., Yu, X., Chen, J., Wang, L., Yang, X., He, S., and Xue, Y. (2020) RIC-seq for global in situ profiling of RNA-RNA spatial interactions. Nature, 582, 432–437.

[45] Lu, Z., Gong, J., and Zhang, Q. C. (2018) PARIS: Psoralen Analysis of RNA Interactions and Structures with High Throughput and Resolution. Methods Mol Biol, 1649, 59–84.

[46] Wu, T., Cheng, A. Y., Zhang, Y., Xu, J., Wu, J., Wen, L., Li, X., Liu, B., Dou, X., Wang, P., Zhang, L., Fei, J., Li, J., Ouyang, Z., and He, C. (2024) KARR-seq reveals cellular higher-order RNA structures and RNA-RNA interactions. Nat Biotechnol, 42, 1909–1920.

[47] Rivas, E., Clements, J., and Eddy, S. R. (2017) A statistical test for conserved RNA struc-ture shows lack of evidence for structure in lncRNAs. Nat Methods, 14, 45–48.

[48] Jin, H., Zhang, C., Zwahlen, M., von Feilitzen, K., Karlsson, M., Shi, M., Yuan, M., Song, X., Li, X., Yang, H., Turkez, H., Fagerberg, L., Uhlén, M., and Mardinoglu, A. (2023) Systematic transcriptional analysis of human cell lines for gene expression landscape and tumor representation. Nat Commun, 14, 5417.

[49] Kalinina, M., Skvortsov, D., Kalmykova, S., Ivanov, T., Dontsova, O., and Pervouchine, D. D. (2021) Multiple competing RNA structures dynamically control alternative splicing in the human ATE1 gene. Nucleic Acids Res, 49, 479–490.

[50] Lovci, M. T., Ghanem, D., Marr, H., Arnold, J., Gee, S., Parra, M., Liang, T. Y., Stark, T. J., Gehman, L. T., Hoon, S., Massirer, K. B., Pratt, G. A., Black, D. L., Gray, J. W., Conboy, J. G., and Yeo, G. W. (2013) Rbfox proteins regulate alternative mRNA splicing through evolutionarily conserved RNA bridges. Nat Struct Mol Biol, 20, 1434–42.

[51] Driscoll, S., Merkuri, F., Chain, F. J. J., and Fish, J. L. (2024) Splicing is dynamically regulated during limb development. Sci Rep, 14, 19944.

[52] Tian, G. G., Li, J., and Wu, J. (2020) Alternative splicing signatures in preimplantation embryo development. Cell Biosci, 10, 33.

[53] Abascal, F., Tress, M. L., and Valencia, A. (2015) The evolutionary fate of alternatively spliced homologous exons after gene duplication. Genome Biol Evol, 7, 1392–403.

[54] Yang, Y., Sun, F., Wang, X., Yue, Y., Wang, W., Zhang, W., Zhan, L., Tian, N., Shi, F., and Jin, Y. (2012) Conservation and regulation of alternative splicing by dynamic inter- and intra-intron base pairings in Lepidoptera 14-3-3ζ pre-mRNAs. RNA Biol, 9, 691–700.

[55] Clouet d’Orval, B., d’Aubenton Carafa, Y., Sirand-Pugnet, P., Gallego, M., Brody, E., and Marie, J. (1991) RNA secondary structure repression of a muscle-specific exon in HeLa cell nuclear extracts. Science, 252, 1823–8.

[56] Pervouchine, D. D. (2018) Towards Long-Range RNA Structure Prediction in Eukaryotic Genes. Genes (Basel), 9, 302.

[57] Conboy, J. G. (2017) Developmental regulation of RNA processing by Rbfox proteins. Wiley Interdiscip Rev RNA, 8, 10.1002/wrna.1398.

[58] Gehman, L. T., Meera, P., Stoilov, P., Shiue, L., O’Brien, J. E., Meisler, M. H., Ares, M., Otis, T. S., and Black, D. L. (2012) The splicing regulator Rbfox2 is required for both cerebellar development and mature motor function. Genes Dev, 26, 445–60.

[59] Gehman, L. T., Stoilov, P., Maguire, J., Damianov, A., Lin, C.-H., Shiue, L., Ares, M., Mody, I., and Black, D. L. (2011) The splicing regulator Rbfox1 (A2BP1) controls neuronal excitation in the mammalian brain. Nat Genet, 43, 706–11.

[60] Singh, R. K., Kolonin, A. M., Fiorotto, M. L., and Cooper, T. A. (2018) Rbfox-Splicing Factors Maintain Skeletal Muscle Mass by Regulating Calpain3 and Proteostasis. Cell Rep, 24, 197–208.

[61] Singh, R. K., Xia, Z., Bland, C. S., Kalsotra, A., Scavuzzo, M. A., Curk, T., Ule, J., Li, W., and Cooper, T. A. (2014) Rbfox2-coordinated alternative splicing of Mef2d and Rock2 controls myoblast fusion during myogenesis. Mol Cell, 55, 592–603.

[62] Jacko, M., Weyn-Vanhentenryck, S. M., Smerdon, J. W., Yan, R., Feng, H., Williams, D. J., Pai, J., Xu, K., Wichterle, H., and Zhang, C. (2018) Rbfox Splicing Factors Promote Neuronal Maturation and Axon Initial Segment Assembly. Neuron, 97, 853–868.e6.

[63] Misra, C., Bangru, S., Lin, F., Lam, K., Koenig, S. N., Lubbers, E. R., Hedhli, J., Murphy, N. P., Parker, D. J., Dobrucki, L. W., Cooper, T. A., Tajkhorshid, E., Mohler, P. J., and Kalsotra, A. (2020) Aberrant Expression of a Non-muscle RBFOX2 Isoform Triggers Cardiac Conduction Defects in Myotonic Dystrophy. Dev Cell, 52, 748–763.e6.

[64] Llorian, M., Schwartz, S., Clark, T. A., Hollander, D., Tan, L.-Y., Spellman, R., Gordon, A., Schweitzer, A. C., de la Grange, P., Ast, G., and Smith, C. W. J. (2010) Position-dependent alternative splicing activity revealed by global profiling of alternative splicing events regulated by PTB. Nat Struct Mol Biol, 17, 1114–23.

[65] Mironov, A., Petrova, M., Margasyuk, S., Vlasenok, M., Mironov, A. A., Skvortsov, D., and Pervouchine, D. D. (2023) Tissue-specific regulation of gene expression via unproductive splicing. Nucleic Acids Res, 51, 3055–3066.

[66] Zuker, M. (2003) Mfold web server for nucleic acid folding and hybridization prediction. Nucleic Acids Res, 31, 3406–15.

[67] Camacho, C., Coulouris, G., Avagyan, V., Ma, N., Papadopoulos, J., Bealer, K., and Madden, T. L. (2009) BLAST+: architecture and applications. BMC Bioinformatics, 10, 421.

[68] Katoh, K. and Standley, D. M. (2013) MAFFT multiple sequence alignment software version 7: improvements in performance and usability. Mol Biol Evol, 30, 772–80.

[69] Edgar, R. C. (2022) Muscle5: High-accuracy alignment ensembles enable unbiased as-sessments of sequence homology and phylogeny. Nat Commun, 13, 6968.

[70] Minh, B. Q., Schmidt, H. A., Chernomor, O., Schrempf, D., Woodhams, M. D., von Haeseler, A., and Lanfear, R. (2020) IQ-TREE 2: New Models and Efficient Methods for Phylogenetic Inference in the Genomic Era. Mol Biol Evol, 37, 1530–1534.

[71] Lorenz, R., Bernhart, S. H., Honer Zu Siederdissen, C., Tafer, H., Flamm, C., Stadler, P. F., and Hofacker, I. L. (2011) ViennaRNA Package 2.0. Algorithms Mol Biol, 6, 26.

[72] Consortium, G. (2013) The Genotype-Tissue Expression (GTEx) project. Nat Genet, 45, 580–5.

[73] Network, C. G. A. R., Weinstein, J. N., Collisson, E. A., Mills, G. B., Shaw, K. R. M., Ozenberger, B. A., Ellrott, K., Shmulevich, I., Sander, C., and Stuart, J. M. (2013) The Cancer Genome Atlas Pan-Cancer analysis project. Nat Genet, 45, 1113–20.

[74] Margasyuk, S., Kuznetsova, A., Zavileyskiy, L., Vlasenok, M., Skvortsov, D., and Pervou-chine, D. D. (2024) Human introns contain conserved tissue-specific cryptic poison exons. NAR Genom Bioinform, 6, lqae163.

[75] Sinha, I. R., Ye, Y., Li, Y., Sandal, P. S., Wong, P. C., Sun, S., and Ling, J. P. (2025) Inhibition of nonsense-mediated decay in TDP-43 deficient neurons reveals novel cryptic exons. bioRxiv, p. 2025.06.28.661837.

[76] Dobin, A., Davis, C. A., Schlesinger, F., Drenkow, J., Zaleski, C., Jha, S., Batut, P., Chais-son, M., and Gingeras, T. R. (2013) STAR: ultrafast universal RNA-seq aligner. Bioinfor-matics, 29, 15–21.

[77] Pervouchine, D. D., Knowles, D. G., and Guigo, R. (2013) Intron-centric estimation of alternative splicing from RNA-seq data. Bioinformatics, 29, 273–4.

[78] Margasyuk, S., Kalinina, M., Petrova, M., Skvortsov, D., Cao, C., and Pervouchine, D. D. (2023) RNA in situ conformation sequencing reveals novel long-range RNA structures with impact on splicing. RNA, 29, 1423–1436.

[79] Margasyuk, S., Zavileyskiy, L., Cao, C., and Pervouchine, D. (2023) Long-range RNA structures in the human transcriptome beyond evolutionarily conserved regions. PeerJ, 11, e16414.

[80] Chernyavskaya, E. SCNa splicing: 10.5281/zenodo.17858476. (December, 2025).

