## Supplementary material for "Ancestral intronic splicing regulatory elements in the SCNα gene family"

### SUPPLEMENTARY INFORMATION: Ancestral intronic splicing regulatory elements in the SCN$\alpha$ gene family

December 17, 2025

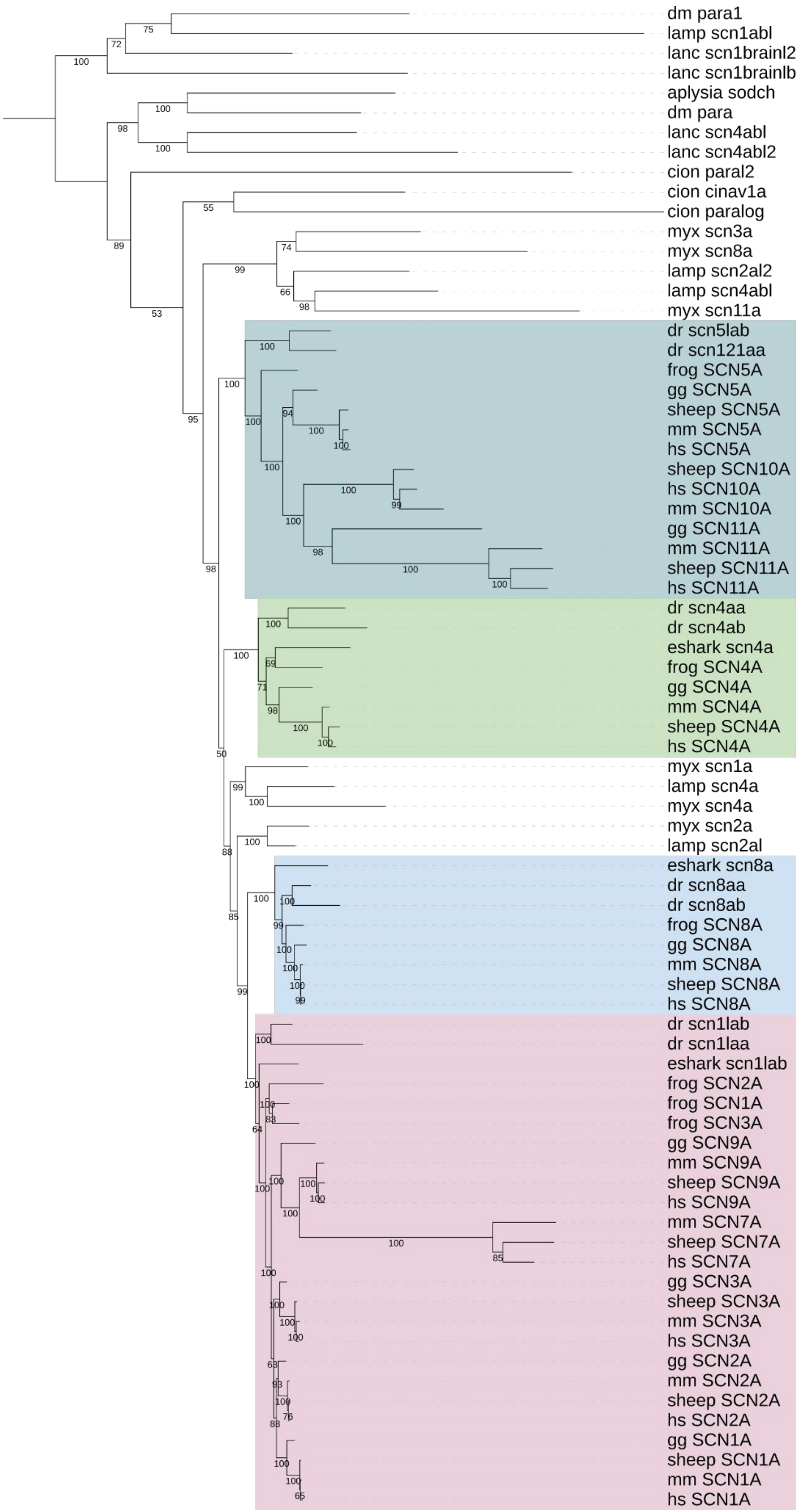

**Figure S1:** Phylogenetic tree of SCN$\alpha$ genes.

**A**

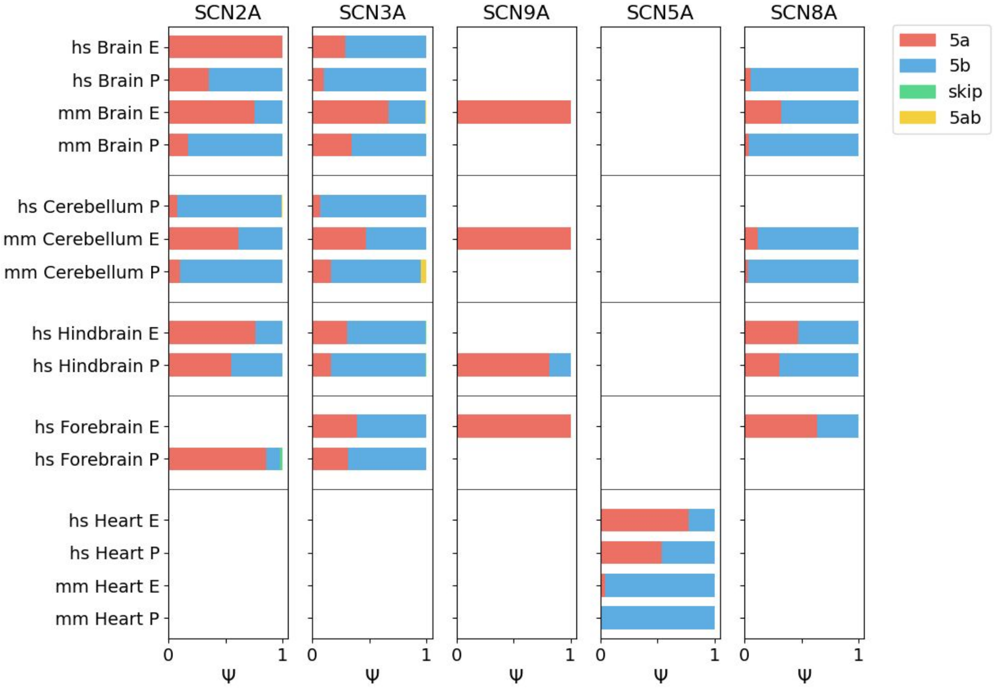

**B**

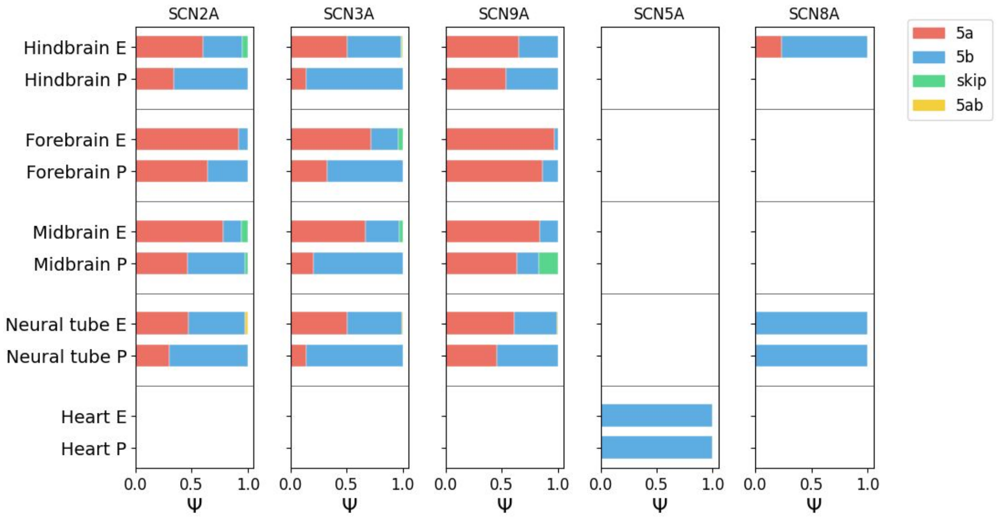

**Figure S2:** Developmental splicing patterns in SCN$\alpha$ genes. The median $\Psi$ value of each isoform from each tissue is shown. **(A)** Comparison of developmental stages between human and mouse across tissues. **(B)** Comparison of different mouse developmental stages by tissue. (E — embryonic, P — postnatal)

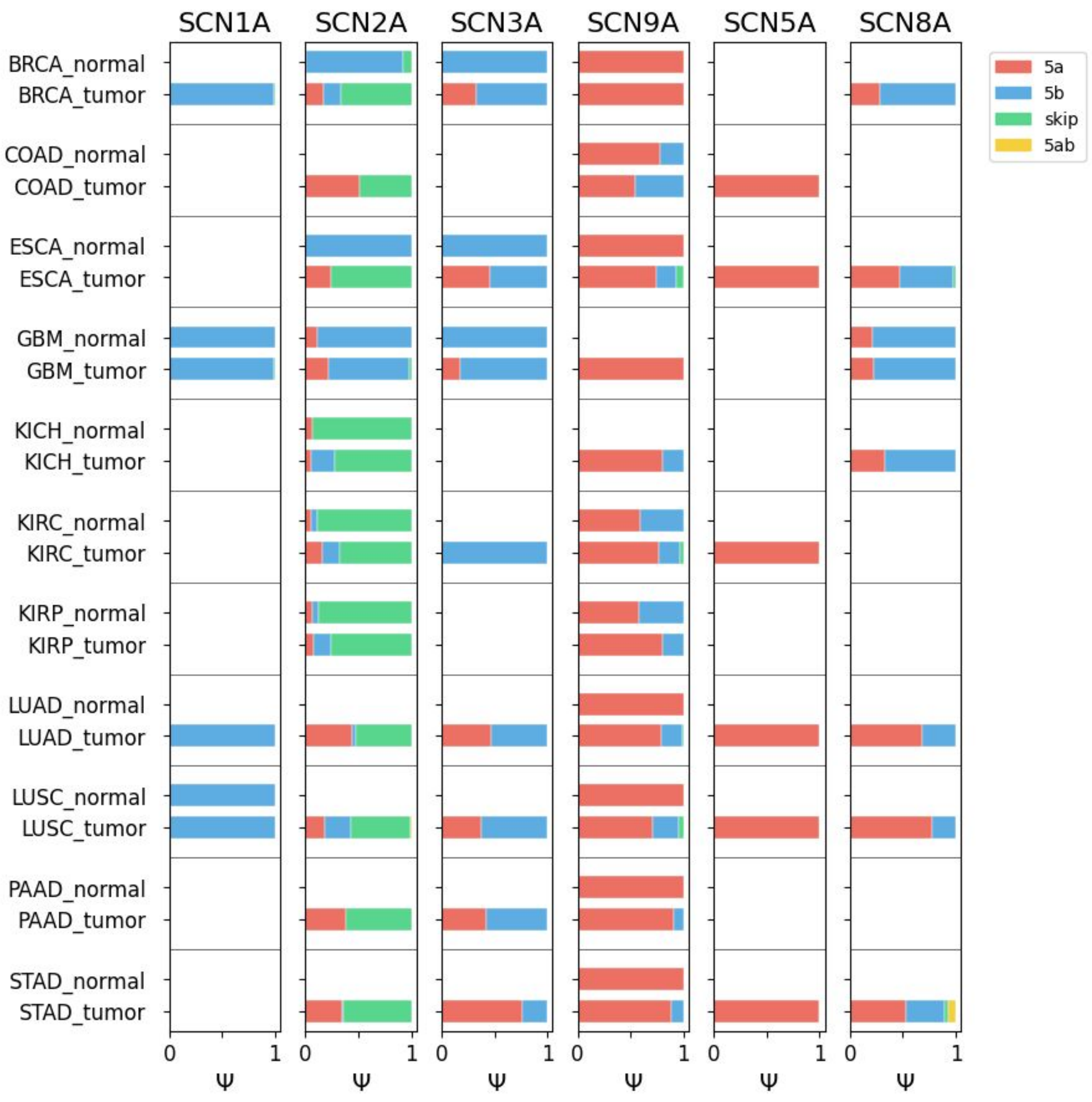

**Figure S3:** Splicing patterns in SCN$\alpha$ genes across TCGA cohorts.

**A**

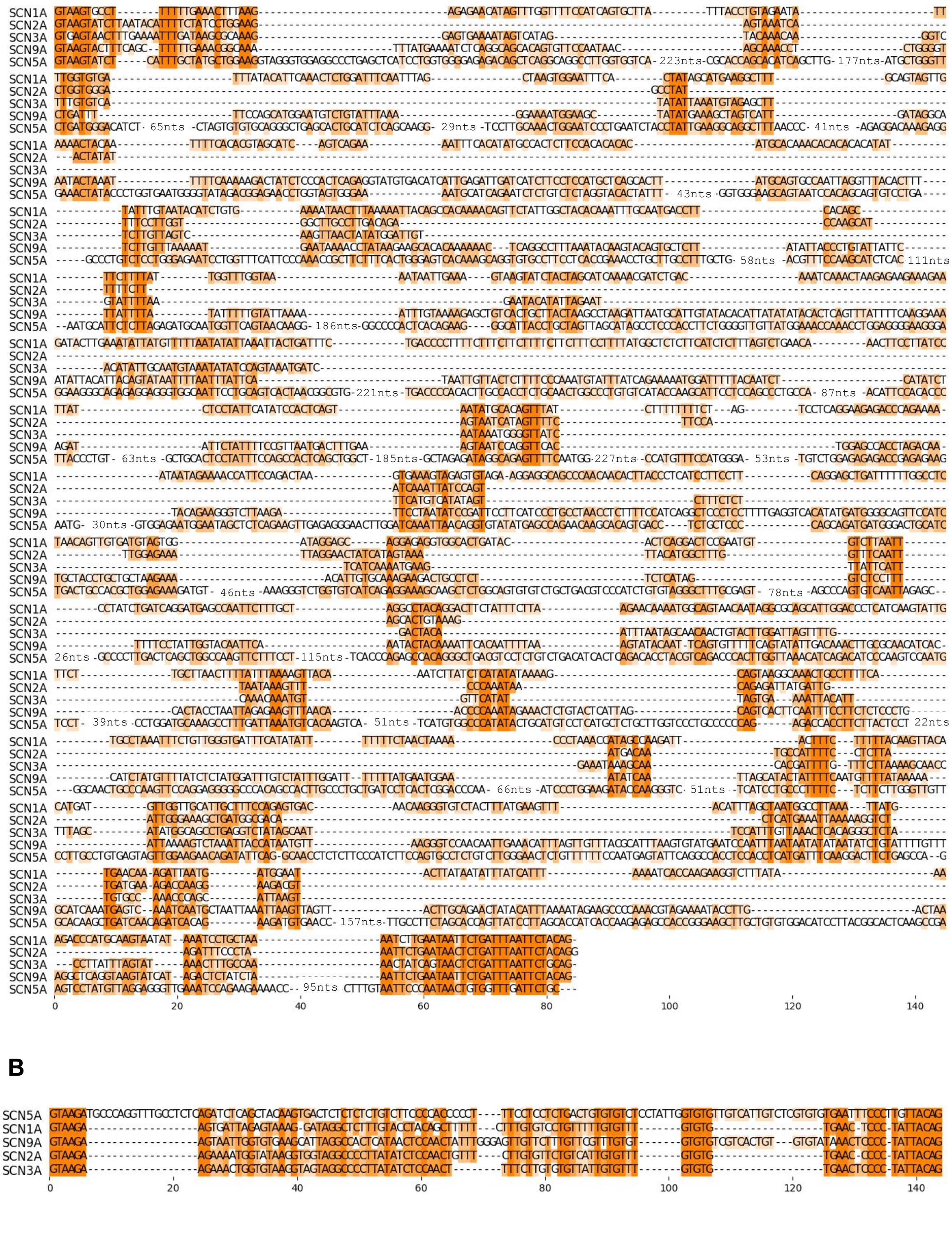

**B**

**Figure S4:** Multiple alignments of intronic sequences of the human *SCN1A*, *SCN2A*, *SCN3A*, *SCN9A*, and *SCN5A* genes. **(A)** The intron between exons 4 and 5a. **(B)** The intron between exons 5a and 5b.

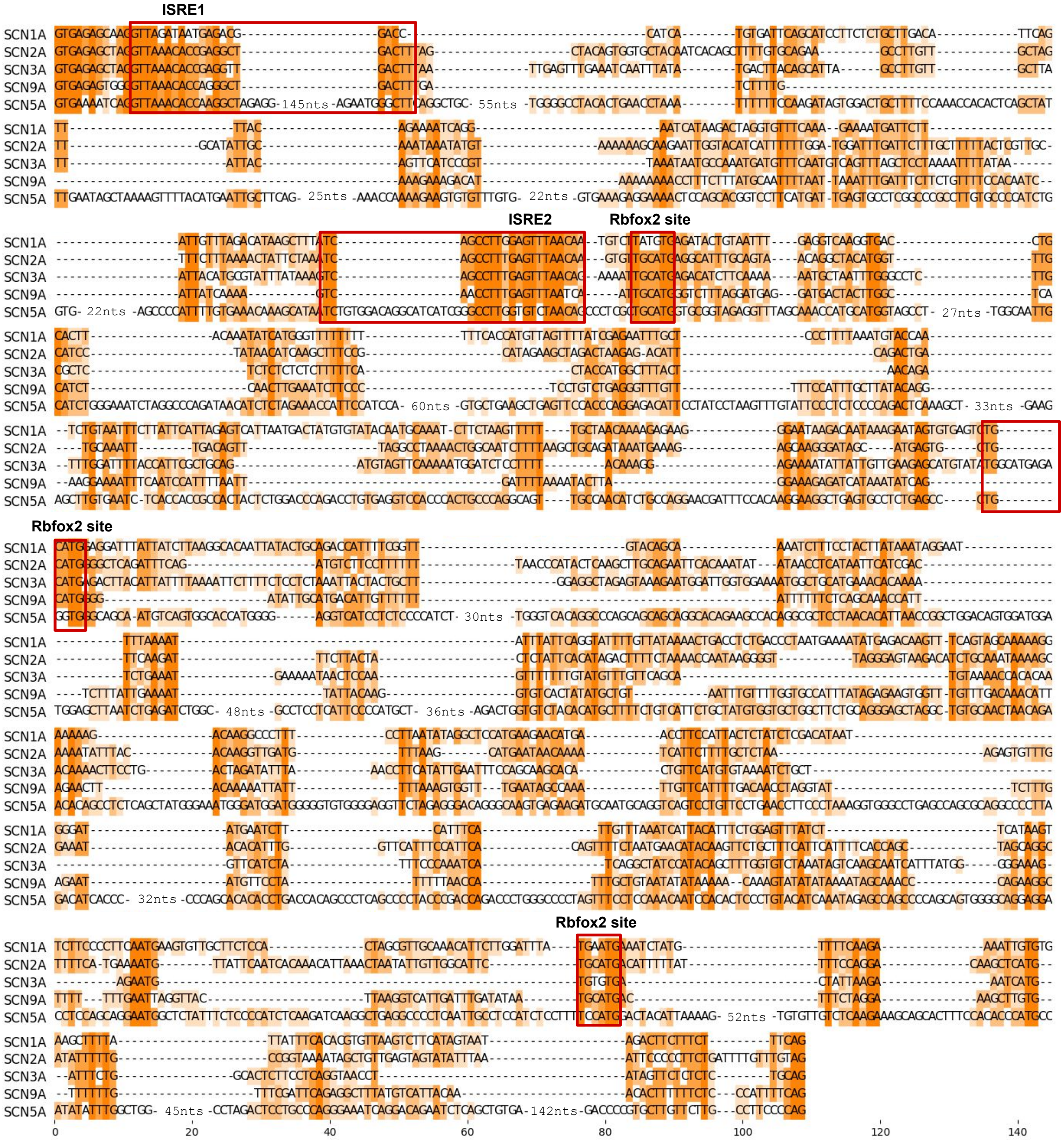

**Figure S5:** A multiple alignment of the intronic sequence between exons 5b and 6 in the human *SCN1A*, *SCN2A*, *SCN3A*, *SCN9A*, and *SCN5A* genes. The ISRE1 and ISRE2 sequences and putative RBFOX2 binding sites are highlighted by red frames.

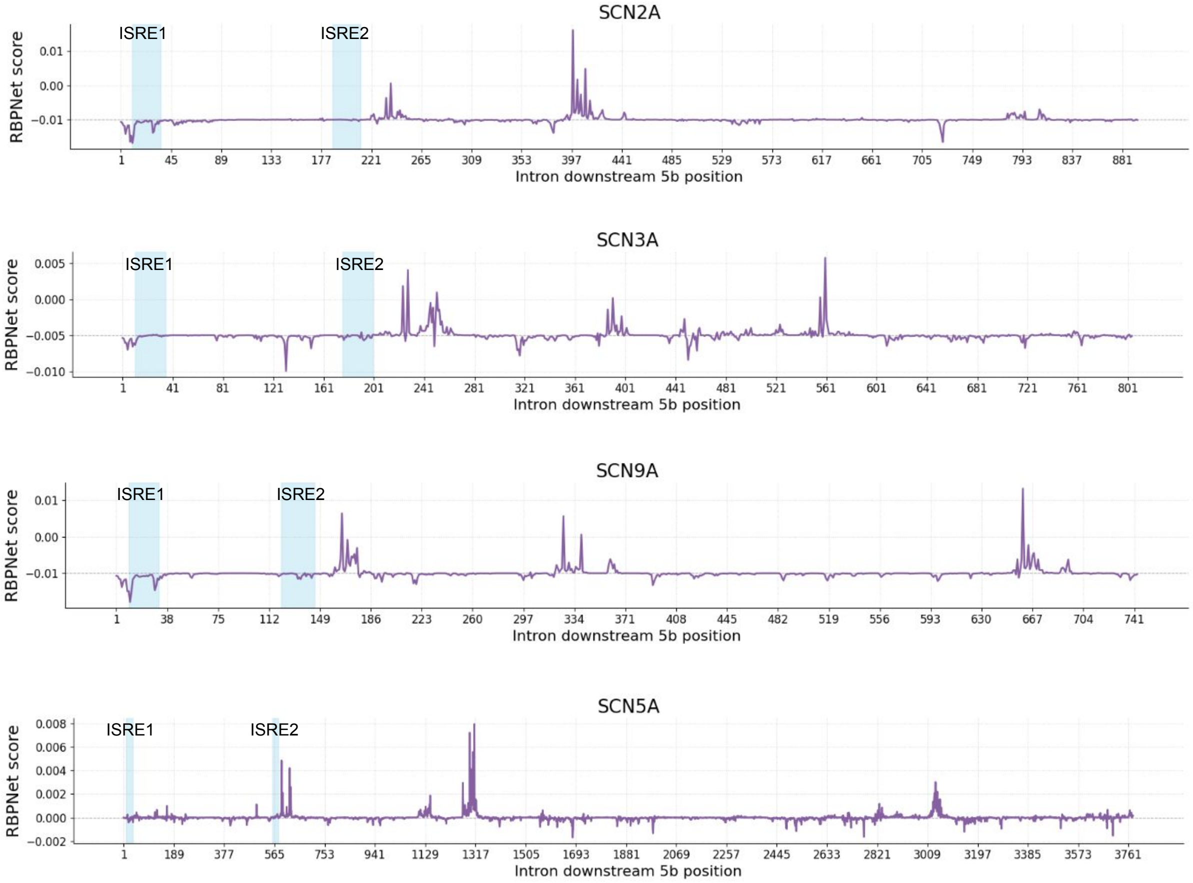

**Figure S6:** RBPNet target score of Rbfox2 binding between exons 5b and 6 in the human *SCN2A*, *SCN3A*, *SCN9A*, and *SCN5A* genes. The locations of ISRE1 and ISRE2 sequences are highlighted blue. Shown in the $y$-axis is the difference between protein-specific target signal and the background control signal.

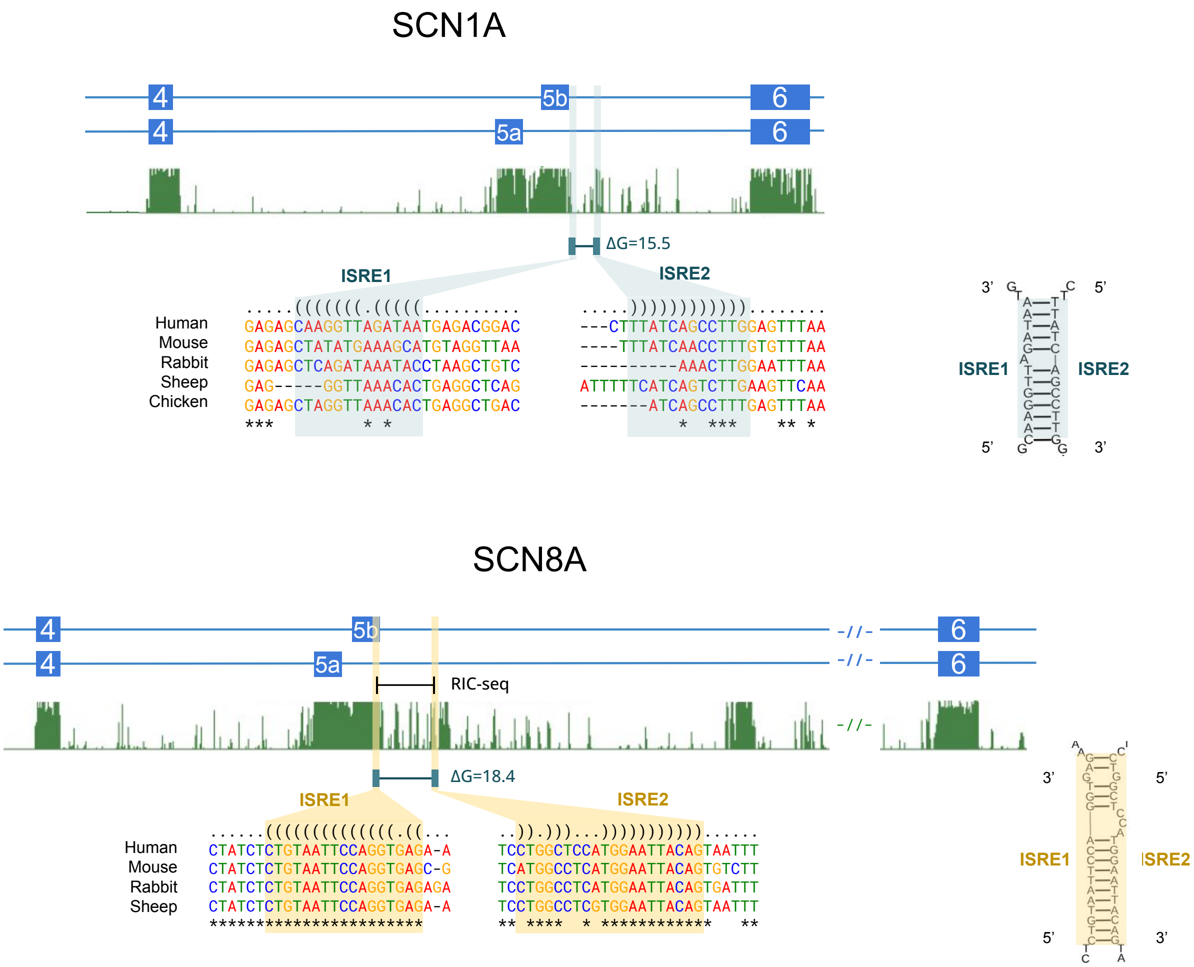

**Figure S7:** The ISRE1 and ISRE2 sequences in *SCN1A* and *SCN8A* genes. The legend is as in Figure 4.

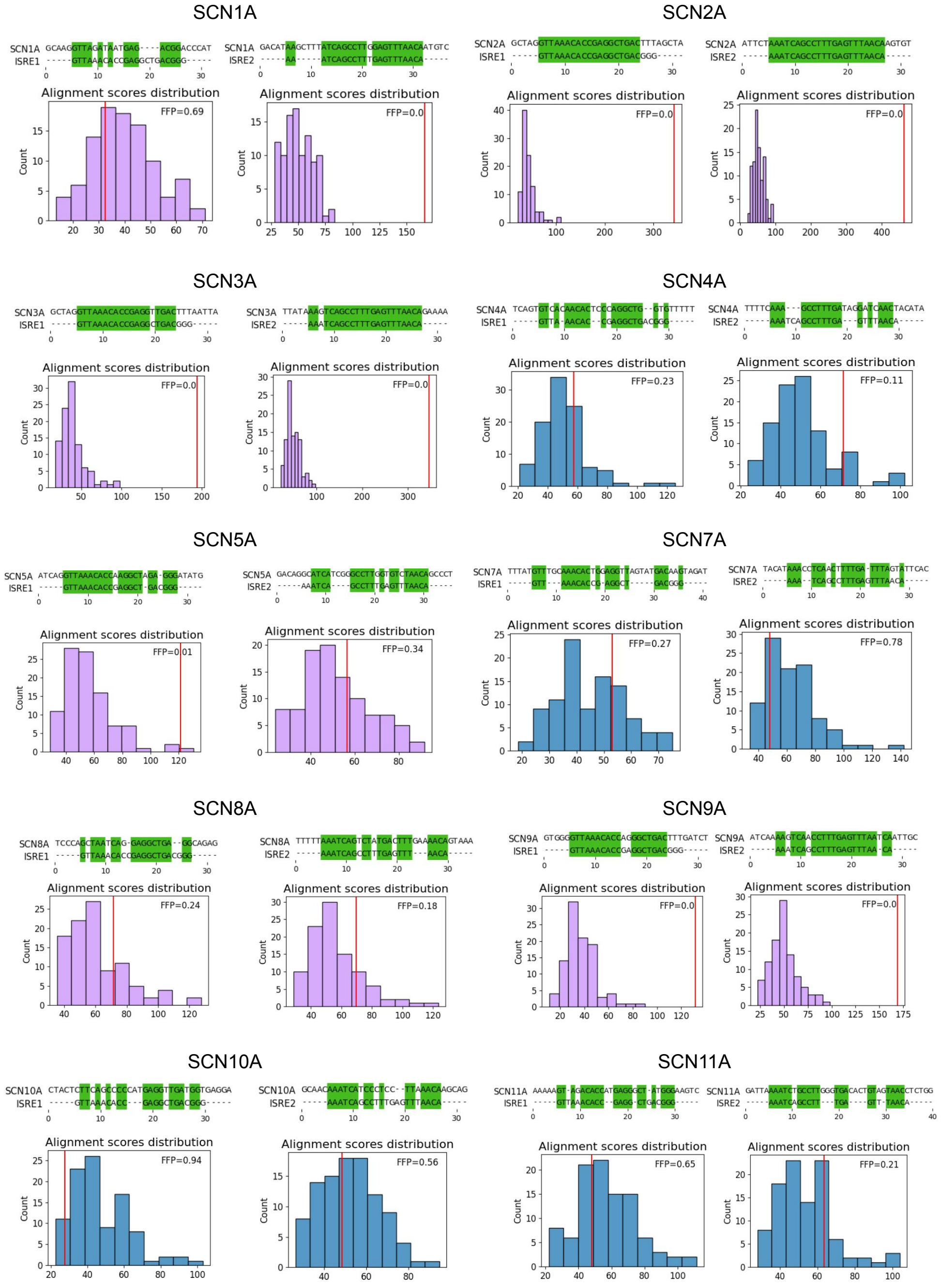

**Figure S8:** Each of the ten panels shows the alignment of the consensus ISRE1 and ISRE2 sequences (Figure 4) to the respective introns between exons 5b and 6 in human SCN$\alpha$ genes. The score of the actual alignment is shown by the red line in histograms below; the histograms represent the distribution of the alignment scores for shuffled intronic sequences (100 replicates). FFP denotes the fraction of shuffled intronic sequences in which the alignment score was greater than the actual score. Purple color denotes SCN$\alpha$ genes with exon 5 duplications. At least one of the two ISRE was found in *SCN1A*, *SCN2A*, *SCN3A*, *SCN5A*, and *SCN9A*.

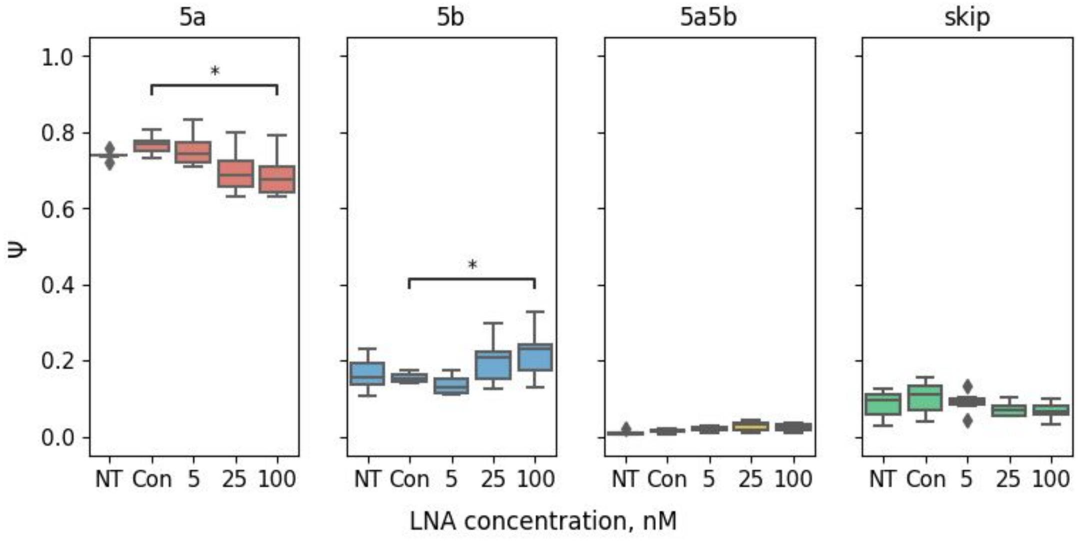

**Figure S9:** In *SCN9A*, ASO2 treatment significantly reduces 5a inclusion and increases 5b inclusion without significant upregulation of non-MXE isoforms (5a5b and skip). NT (nontreated control); Con (control ASO). Asterisks (*) denote statistically significant differences at the 5

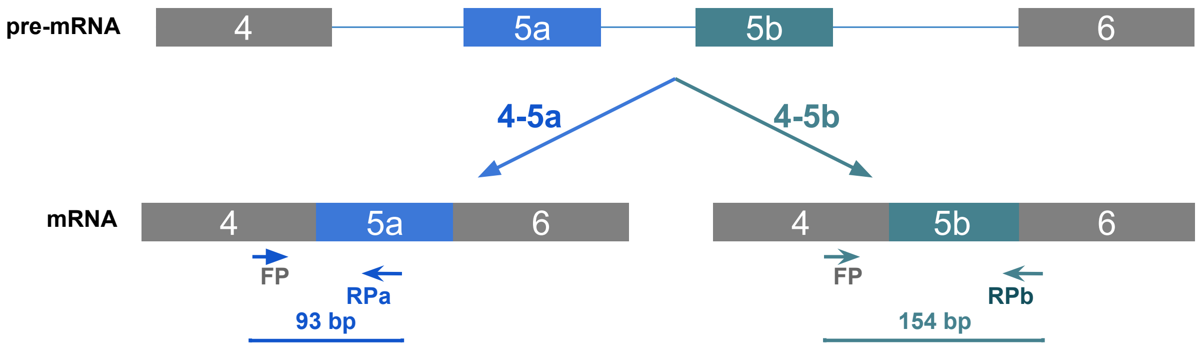

**Figure S10:** RT-PCR primer locations in *SCN9A* exons 5a and 5b. Since exons 5a and 5b in *SCN9A* are both 92-nts-long, two different reverse primers were used to visualize them in the gel. FP denotes the common forward primer; RPa and RPb denote reverse primers complementary to exons 5a and 5b, respectively.

| Species | Gene | Closest human homolog | Specie’s exon 5 identity | Human exon 5b identity | Assembly and coordinates |
|---|---|---|---|---|---|
| *M. marmota* | SCN1A | SCN1A | 56% | 59% | marMar2.1 CZRN01000006.1:53124421-53124513 |
| *O. aries* | SCN2A | SCN2A | 97% | 100% | Oar_v3.1 2:143095931-143096021 |
| *X. tropicalis* | SCN2A | SCN2A | 47% | 47% | UCB_Xtro_10.0 9:70322548-70322640 |
| *D. rerio* | SCN1A | SCN1A | 97% | 97% | GRCz11 6:10299249-10299341 |
| *C. milii* | SCN1A | SCN1A | 67% | 65% | Cm-6.1.3 KI635868.1:1195059-1195151 |
| *B. lanceolatum* | LOC136422135 | SCN4A | 96% | 76% | klBraLanc5.hap2 16:982220-982312 |
| *C. intestinalis* | LOC100180733 | SCN3A | 50% | 43% | KH 9:4025436-4025528 |

**Table S1:** Unannotated exon duplication events. The columns 4 and 5 show the percentage of identical amino acids in the alignment.

| Gene | Huh7 (nTPM) | A549 (nTPM) |
|---|---|---|
| SCN5A | 0 | 0.4 |
| SCN8A | 1.5 | 1.7 |
| SCN9A | 32.5 | 6.1 |

**Table S2:** nTPM expression levels (transcripts per million) for SCN$\alpha$ genes where in at least one cell line (Huh7 or A549) the expression level was above zero.

| Species | | Abbr. | Gene | Ensembl / NCBI ID |
|---|---|---|---|---|
| *H. sapiens* | Human | hs | SCN1A | ENSG00000144285 |
| | | hs | SCN2A | ENSG00000136531 |
| | | hs | SCN3A | ENSG00000153253 |
| | | hs | SCN9A | ENSG00000169432 |
| | | hs | SCN7A | ENSG00000136546 |
| | | hs | SCN5A | ENSG00000183873 |
| | | hs | SCN10A | ENSG00000185313 |
| | | hs | SCN11A | ENSG00000168356 |
| | | hs | SCN8A | ENSG00000196876 |
| | | hs | SCN4A | ENSG00000007314 |
| *M. musculus* | Mouse | mm | SCN1A | ENSMUSG00000064329 |
| | | mm | SCN2A | ENSMUSG00000075318 |
| | | mm | SCN3A | ENSMUSG00000057182 |
| | | mm | SCN9A | ENSMUSG00000075316 |
| | | mm | SCN7A | ENSMUSG00000034810 |
| | | mm | SCN5A | ENSMUSG00000032511 |
| | | mm | SCN10A | ENSMUSG00000034533 |
| | | mm | SCN11A | ENSMUSG00000034115 |
| | | mm | SCN8A | ENSMUSG00000023033 |
| | | mm | SCN4A | ENSMUSG00000001027 |
| *O. aries* | Sheep | sheep | SCN1A | ENSOARG00020010370 |
| | | sheep | SCN2A | ENSOARG00020007222 |
| | | sheep | SCN3A | ENSOARG00020007511 |
| | | sheep | SCN9A | ENSOARG00020006902 |
| | | sheep | SCN7A | ENSOARG00020035820 |
| | | sheep | SCN5A | ENSOARG00020010796 |
| | | sheep | SCN10A | ENSOARG00020016009 |

Continued on next page

**Table S3 – continued from previous page**

| Species | | Abbr. | Gene | Ensembl / NCBI ID |
|---|---|---|---|---|
| | | sheep | SCN11A | ENSOARG00020016401 |
| | | sheep | SCN8A | ENSOARG00020015921 |
| | | sheep | SCN4A | ENSOARG00020016096 |
| *G. gallus* | Chicken | gg | SCN1A | ENSGALG00010026770 |
| | | gg | SCN2A | ENSGALG00010025597 |
| | | gg | SCN3A | ENSGALG00010025628 |
| | | gg | SCN9A | ENSGALG00010025505 |
| | | gg | SCN5A | ENSGALG00010027095 |
| | | gg | SCN11A | ENSGALG00010026737 |
| | | gg | SCN8A | ENSGALG00010022264 |
| | | gg | SCN4A | ENSGALG00010023635 |
| *X. tropicalis* | Frog | frog | SCN1A | ENSXETG00000020846 |
| | | frog | SCN2A | ENSXETG00000021004 |
| | | frog | SCN3A | ENSXETG00000008965 |
| | | frog | SCN5A | ENSXETG00000004251 |
| | | frog | SCN8A | ENSXETG00000008958 |
| | | frog | SCN4A | ENSXETG00000014235 |
| *D. rerio* | Zebrafish | dr | scn1aa | ENSDARG00000086819 |
| | | dr | scn1lab | ENSDARG00000062744 |
| | | dr | scn5lab | ENSDARG00000102312 |
| | | dr | scn12aa | ENSDARG00000090724 |
| | | dr | scn8aa | ENSDARG00000005775 |
| | | dr | scn8ab | ENSDARG00000112543 |
| | | dr | scn4aa | ENSDARG00000098738 |
| | | dr | scn4ab | ENSDARG00000034588 |
| *C. milii* | Elephant shark | eshark | scn1lab | ENSCMIG00000008242 |

Continued on next page

**Table S3 – continued from previous page**

| Species | | Abbr. | Gene | Ensembl / NCBI ID |
|---|---|---|---|---|
| | | eshark | scn8a | ENSCMIG00000018145 |
| | | eshark | scn4a | ENSCMIG00000008660 |
| *P. marinus* | Lamprey | lamp | scn1abl | LOC116942140 |
| | | lamp | scn2al | LOC116954695 |
| | | lamp | scn4a | LOC103091816 |
| | | lamp | scn2al2 | LOC116940894 |
| | | lamp | scn4abl | LOC116948895 |
| *M. glutinosa* | Myxine | myx | scn1a | LOC137417442 |
| | | myx | scn2a | LOC137438074 |
| | | myx | scn3a | LOC137429583 |
| | | myx | scn4a | LOC137416790 |
| | | myx | scn8a | LOC137433392 |
| | | myx | scn11a | LOC137433391 |
| *B. lanceolatum* | Lancelet | lanc | scn4abl | LOC136422130 |
| | | lanc | scn4abl2 | LOC136422135 |
| | | lanc | scn1brainlb | LOC136424728 |
| | | lanc | scn1brainl2 | LOC136428574 |
| *C. intestinalis* | Ciona | cion | CiNav1a | ENSCING00000002389 |
| | | cion | paralog1 | ENSCING00000005490 |
| | | cion | paralog2 | ENSCING00000012159 |
| *D. melanogaster* | Drosophila | dm | para | FBgn0285944 |
| | | dm | para1 | FBgn0085434 |
| *A. californica* | Aplysia | aplysia | sodch | AAC47457 |

**Table S3:** Accession numbers of annotated SCN$\alpha$ genes in 13 studied species.

| Tissue | Number of samples |
|---|---|
| Artery - Aorta | 246 |
| Artery - Coronary | 140 |
| Artery - Tibial | 357 |
| Brain - Amygdala | 81 |
| Brain - Anterior cingulate cortex (BA24) | 99 |
| Brain - Caudate (basal ganglia) | 134 |
| Brain - Cerebellar Hemisphere | 115 |
| Brain - Cerebellum | 144 |
| Brain - Cortex | 132 |
| Brain - Frontal Cortex (BA9) | 117 |
| Brain - Hippocampus | 103 |
| Brain - Hypothalamus | 104 |
| Brain - Nucleus accumbens (basal ganglia) | 123 |
| Brain - Putamen (basal ganglia) | 103 |
| Brain - Spinal cord (cervical c-1) | 76 |
| Brain - Substantia nigra | 71 |
| Breast - Mammary Tissue | 218 |
| Colon - Sigmoid | 173 |
| Fallopian Tube | 7 |
| Heart - Atrial Appendage | 218 |
| Heart - Left Ventricle | 267 |
| Lung | 372 |
| Minor Salivary Gland | 70 |
| Muscle - Skeletal | 470 |
| Nerve - Tibial | 334 |
| Ovary | 108 |
| Pancreas | 192 |
| Pituitary | 124 |
| Testis | 198 |

**Table S4:** The number of tissue samples in the GTEx dataset.

| TCGA Cohort | Number of paired samples |
|---|---|
| TCGA-BRCA | 112 |
| TCGA-COAD | 41 |
| TCGA-ESCA | 8 |
| TCGA-KICH | 23 |
| TCGA-KIRC | 72 |
| TCGA-KIRP | 31 |
| TCGA-LUAD | 57 |
| TCGA-LUSC | 49 |
| TCGA-PAAD | 4 |
| TCGA-STAD | 27 |

**Table S5:** The number of tumor and matched normal tissue samples and number of paired tumor-normal samples in 10 TCGA cohorts that were selected for the analysis.

| Tissue | Embryonic | Postnatal |
|---|---|---|
| Forebrain | 6 | 2 |
| Heart | 8 | 2 |
| Hindbrain | 9 | 2 |
| Midbrain | 13 | 2 |
| Neural tube | 8 | 2 |

**Table S6:** The number of embryonic and postnatal samples from the study of mammalian organ development.

| Primer | Sequence |
|---|---|
| scn9a_m1_f | TTTCAgTCgggACCACAAATTgggATCTTTTgAAAgAAAgACATAAAAAAAACC |
| scn9a_m1_r | CCACTCTCACCTggAATgACTg |
| scn9a_m2_r | TgATAATgATTgTggAAAACAgAAgAAATC |
| scn9a_m2_f | CTAATTTgAgTTTCCAACTgAAAAATTgCATgggTCTTTAggATgAgg |

**Table S7:** Mutagenesis primers.

| siRNA target | Sequence |
|---|---|
| RBFOX2_s | 5’-CAGACACAAAGUAGUGAAAdTdT-3’ |
| RBFOX2_as | 5’-UUUCACUACUUUGUGUCUGdTdT-3’ |
| Luc_s | 5’-CUUACGCUGAGUACUUCGAdTdT-3’ |
| Luc_as | 5’-UCGAAGUACUCAGCGUAAGdTdT-3’ |

**Table S8:** siRNA sequences.

| Isoform | Primer | Sequence |
|---|---|---|
| 5a | scn9a_ex4_f | TTCTTCGTGACCCGTGGAAC |
| | scn9a_5a_in_r | ACAATTGTCTTCAGGCCTGGGA |
| 5b | scn9a_5b_in_f | GTCGTCATTGTTTTTGATATGTGACAG |
| | scn9a_qpcr_r | ACAGAACACAGTCAGGATCATG |
| 5a5b | scn9a_ex4_f | TTCTTCGTGACCCGTGGAAC |
| | scn9a_5a5b_r | CAAACTCTGTCACATATCTGGG |
| Skip | scn9a_4-6_in_f | CGTCATTGTTTTTGGCCTGAAG |
| | scn9a_ex6_r | CTTCTTCACTCTCTAGGGTATTCAT |
| Rbfox2-KD | rbfox2_f | ggATTCgggTTCgTAACTTTCg |
| | rbfox2_r | CATTTgCATATggTgTgACCATCT |

**Table S9:** RT-PCR and RT-qPCR primers.
